# The maternal effect gene *Wds* controls *Wolbachia* titer in *Nasonia*

**DOI:** 10.1101/256909

**Authors:** Lisa J. Funkhouser-Jones, Edward J. van Opstal, Ananya Sharma, Seth R. Bordenstein

**Author notes:** These authors contributed equally to this work (Lisa J. Funkhouser-Jones, Edward J. van Opstal). Corresponding authors. (SRB), (LJF), (EjvO).

## Abstract

Maternal transmission of intracellular microbes is pivotal in establishing long-term, intimate symbioses. For germline microbes that exert negative reproductive effects on their hosts, selection can theoretically favor the spread of host genes that counteract the microbe’s harmful effects. Here, we leverage a major difference in bacterial (*Wolbachia pipientis*) titers between closely-related wasp species with forward genetic, transcriptomic, and cytological approaches to map two quantitative trait loci that suppress bacterial titers via a maternal effect. Fine mapping and knockdown experiments identify the gene *Wolbachia density suppressor* (*Wds*), which dominantly suppresses bacterial transmission from mother to embryo. *Wds* evolved by lineage-specific non-synonymous changes driven by positive selection. Collectively, our findings demonstrate that a genetically simple change arose by Darwinian selection in less than a million years to regulate maternally transmitted bacteria via a dominant, maternal effect gene.

## INTRODUCTION

Many animals harbor microorganisms that participate in beneficial processes as diverse as nutritional uptake and metabolism [1, 2], immune cell development [3, 4], and pathogen resistance [5, 6]. However, even innocuous microbes can become harmful when not properly regulated [7, 8]. Moreover, intracellular symbionts that are maternally transmitted over multiple host generations can impose long-term, negative fitness effects on their hosts [9, 10]. In these intimate and enduring symbioses, hosts are predicted to evolve suppression that reduces the harmful effects of the symbiont [11, 12]. However, little is known about the genes and evolutionary forces that underpin regulation of maternally-transmitted symbionts despite the repeated and independent origins of maternal transmission in diverse host taxa [13]. Reverse genetic studies in insects suggest immune or developmental genes may evolve to affect endosymbiont densities [14–16] but, to our knowledge, no studies have utilized forward genetic approaches to identify the gene(s) underlying variation in host regulation of maternally-transmitted symbionts.

Here, we utilize a major host interspecific difference in titers of the maternally-transmitted bacteria *Wolbachia* and quantitative trait loci analyses to identify a maternal effect, suppressor gene in the *Nasonia* model. *Nasonia* (Order: Hymenoptera) is a genus of parasitoid wasps comprised of four closely-related species, with *N. vitripennis* last sharing a common ancestor with the other three species approximately one million years ago [17, 18]. In the lab, interspecific crosses of *Nasonia* species with the same or no *Wolbachia* produce viable and fertile hybrid females, which permits the transfer of genetic or cytoplasmic material (including intracellular *Wolbachia*) between *Nasonia* species. Consequently, *Nasonia* is a powerful model for studying the quantitative genetics of interspecific variation in host traits such as wing size [19, 20], head shape [21], and sex pheromones [22].

*Wolbachia* (Order Rickettsiales) live intracellularly in 40-52% of all arthropod species [23, 24], and they are predominantly transmitted transovarially with occasional transfer between host species on an evolutionary timescale [25, 26]. In most insects, including in *Nasonia*, *Wolbachia* function as reproductive parasites that manipulate host reproduction through a variety of mechanisms to achieve a greater proportion of infected females in the host population [27, 28]. Both efficient maternal transmission and host reproductive manipulation often depend upon sufficiently high within-host *Wolbachia* densities [29, 30]; however, overproliferation of *Wolbachia* can drastically reduce lifespan in *Drosophila* [10], mosquitoes [31, 32] and terrestrial isopods [33]. Thus, coadaptation between arthropod hosts and *Wolbachia* strain(s) promotes genetic and phenotypic changes that impact transmission of *Wolbachia* densities [34–37]. When co-adapted host and symbiont pairs are disrupted through experimental transfer of *Wolbachia* into a naïve host, control of the symbiosis is often lost, leading to overproliferation, expanded tissue tropism of *Wolbachia*, changes in bacteriophage activity, and/or fitness costs not observed in the original host species [33, 37, 38].

Each *Nasonia* species is naturally infected with different *Wolbachia* strains that were mostly acquired through horizontal transfer but in rare cases have since co-diverged with their host wasp species [39]. Introgression of a specific *Wolbachia* strain (*w*VitA) from one *Nasonia* species (*N. vitripennis*) to a naïve, closely-related species (*N. giraulti*) results in a major perturbation in which the relative *Wolbachia* densities increase by two orders of magnitude with an associated reduction in fecundity in *N. giraulti* [37]. Importantly, *w*VitA densities and *Nasonia* fecundity return to normal when *w*VitA is crossed back into a *N. vitripennis* genomic background from the high-density *N. giraulti* line (IntG). Since both the native *N. vitripennis* and the *w*VitA-infected *N. giraulti* IntG lines have the same cytotype, the interspecific *Wolbachia* density variation is established by variation in the host nuclear genome [37]. In this study, we utilize several forward genetic techniques in *Nasonia* to dissect the genetic, evolutionary, and cytological basis of maternal regulation of *Wolbachia*. The varied approaches culminate in the characterization of two quantitative trait loci and the discovery of *Wolbachia density suppressor* (*Wds*), a positively-selected, maternal effect gene that suppresses the transmission of *Wolbachia*.

## RESULTS

### *Nasonia vitripennis* dominantly suppresses *w*VitA titers through a maternal genetic effect

To determine the inheritance pattern of *w*VitA densities, we reciprocally crossed *N. vitripennis* (low-density) and *N. giraulti* IntG (high-density) individuals and measured the *Wolbachia* densities of F1 female hybrids using quantitative PCR (qPCR) for a single-copy *Wolbachia* gene (*groEL*) normalized to a *Nasonia* gene (*NvS6K*) (Figure 1A). The average F1 female pupal *Wolbachia* densities from pure-breeding *N. vitripennis* (N = 5) and *N. giraulti* control families (N = 5) were 0.057 ± 0.004 and 4.805 ± 1.071 (mean ± S.E.M.), respectively, which represents an 84-fold interspecific difference in *Wolbachia* titers and is consistent with previous studies [37]. Interestingly, while F1 hybrid females from both crosses had identical genotypes (i.e., heterozygous at all loci) and the same cytotype (*N. vitripennis*), the average *Wolbachia* densities in reciprocal F1 hybrid females were significantly different at 0.149 ± 0.029 vs 1.746 ± 0.187 (Figure 1A, N = 10 for both crosses, Kruskal-Wallis test: H = 24.99, df = 3, p < 0.0001, Dunn’s multiple comparisons test: p = 0.03).

**Figure 1.**
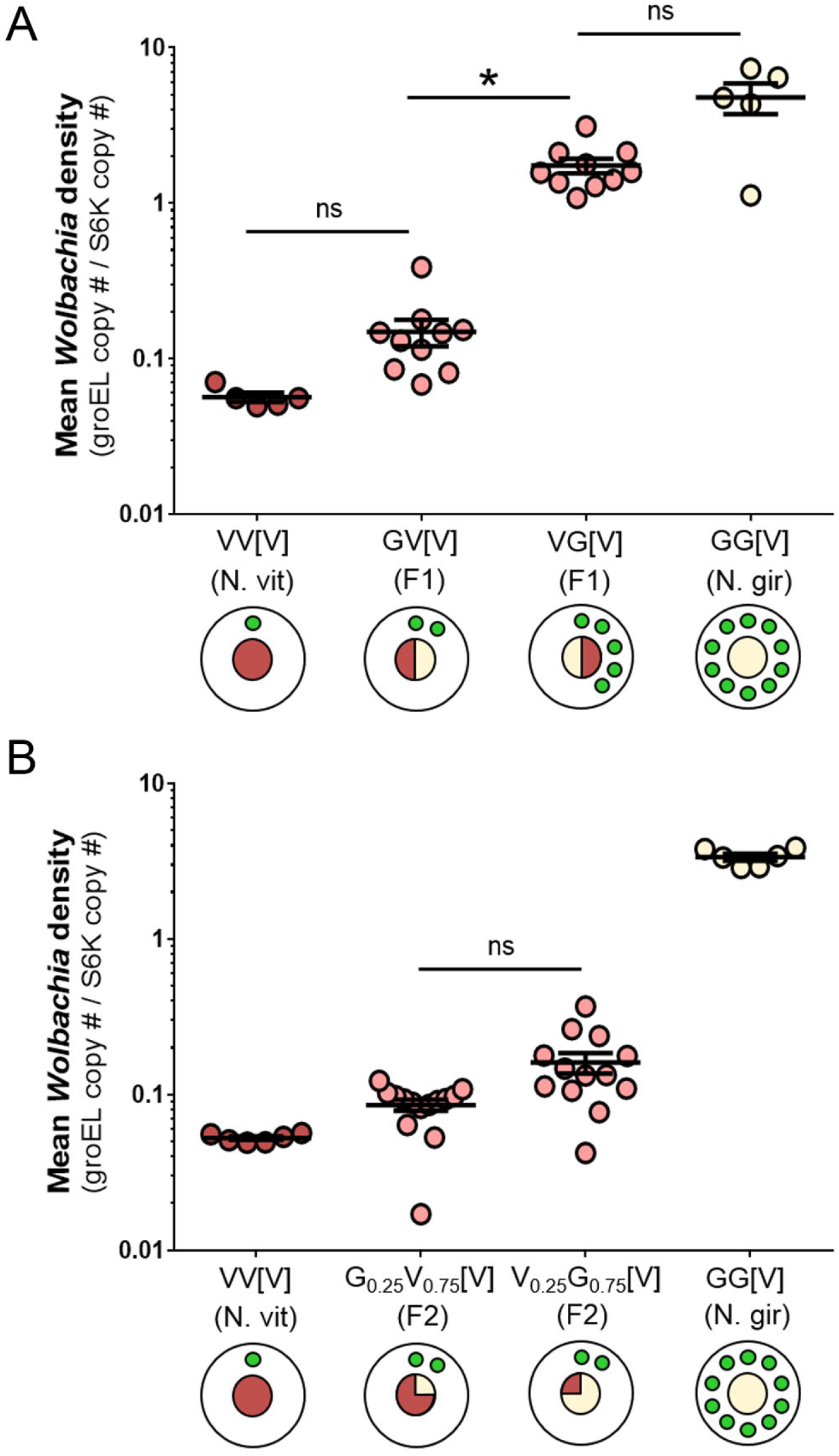
Disparities in *w*VitA titers are established through a maternal genetic effect. (A) *w*VitA densities in female pupae from parental *N. vitripennis* 12.1 and *N. giraulti* IntG, and their reciprocal F1 hybrids. (B) *w*VitA densities in F2 pupae from F1 females backcrossed to their paternal line. For each cross, genotype (male x female) is followed by cytotype in brackets (V = *N. vitripennis*, G = *N. giraulti*, number = estimated proportion of genotype). For the circle diagrams, the inner circle represents the expected percentage of the nuclear genome that is of *N. vitripennis* (red) or *N. giraulti* (cream) origin. Green circles represent *w*VitA load (not drawn to scale). Error bars represent mean ± S.E.M. *p < 0.05, post-hoc Dunn’s test.

To test whether the difference in F1 *Wolbachia* densities was due to maternal *Wolbachia* load or to a maternal genetic effect, we backcrossed F1 females to their paternal line and pooled five female F2 pupae per F1 mother for qPCR (Figure 1B). If a maternal genetic effect regulates *Wolbachia* densities, F2 pupae from both experimental lines should have similar *Wolbachia* levels since F1 hybrid mothers are genotypically identical. Indeed, the densities of F2 pupal offspring of both low- and high-density F1 mothers were not significantly different (Figure 1B, 0.086 ± 0.007, N = 14, and 0.161 ± 0.024, N = 13, respectively, Dunn’s multiple comparisons test: p = 0.18), supporting the inference that maternal nuclear genotype plays an important role in regulating *Wolbachia* densities. Furthermore, since the densities of both F2 hybrid groups were more similar to the *N. vitripennis* control (0.053 ± 0.001, N = 6) than to the *N. giraulti* control (3.364 ± 0.174, N = 6), the *N. vitripennis* suppression gene(s) producing the low *Wolbachia* density phenotype is dominant (Figure 1B).

### Disparities in embryonic *w*VitA levels between *N. vitripennis* and *N. giraulti* are established during oogenesis

We previously showed that variation in *w*VitA loads between *N. vitripennis* and *N. giraulti* exists in early embryos, with strict posterior localization of *w*VitA in *N. vitripennis* that is perturbed in *N. giraulti* embryos [37]. Since the *w*VitA density disparity is partially controlled through a maternal genetic effect (Figure 1B), we reasoned that an embryonic disparity in *w*VitA densities between *N. vitripennis* and *N. giraulti* is likely established in the egg chamber during oogenesis (Figure 2A). Indeed, nucleic acid staining with SYTOX Green revealed that fewer *w*VitA bacteria are present in stage three *N. vitripennis* egg chambers (Figure 2B) than in *N. giraulti* ones at the same stage (Figure 2E, Figure S1). Once in the oocyte, *w*VitA localizes to the posterior pole at low density in *N. vitripennis* (Figure 2B) but occurs at high density in the posterior pole with an expanded distribution toward the anterior end of the oocyte in *N. giraulti* (Figure 2E). The lack of puncta in ovaries from uninfected *N. vitripennis* confirms that the SYTOX nucleic acid dye effectively stains *Wolbachia* but no other cytoplasmic elements, such as mitochondria (Figure 2D). Altogether, the distribution and density of *w*VitA in oocytes of these two *Nasonia* species perfectly mirror their embryonic *w*VitA patterns (Figure 2C and 2F), suggesting that *Wolbachia* density differences are mainly established during oogenesis and regulated through maternal effect genes, rather than zygotic genes expressed later in embryonic development.

**Figure 2.**
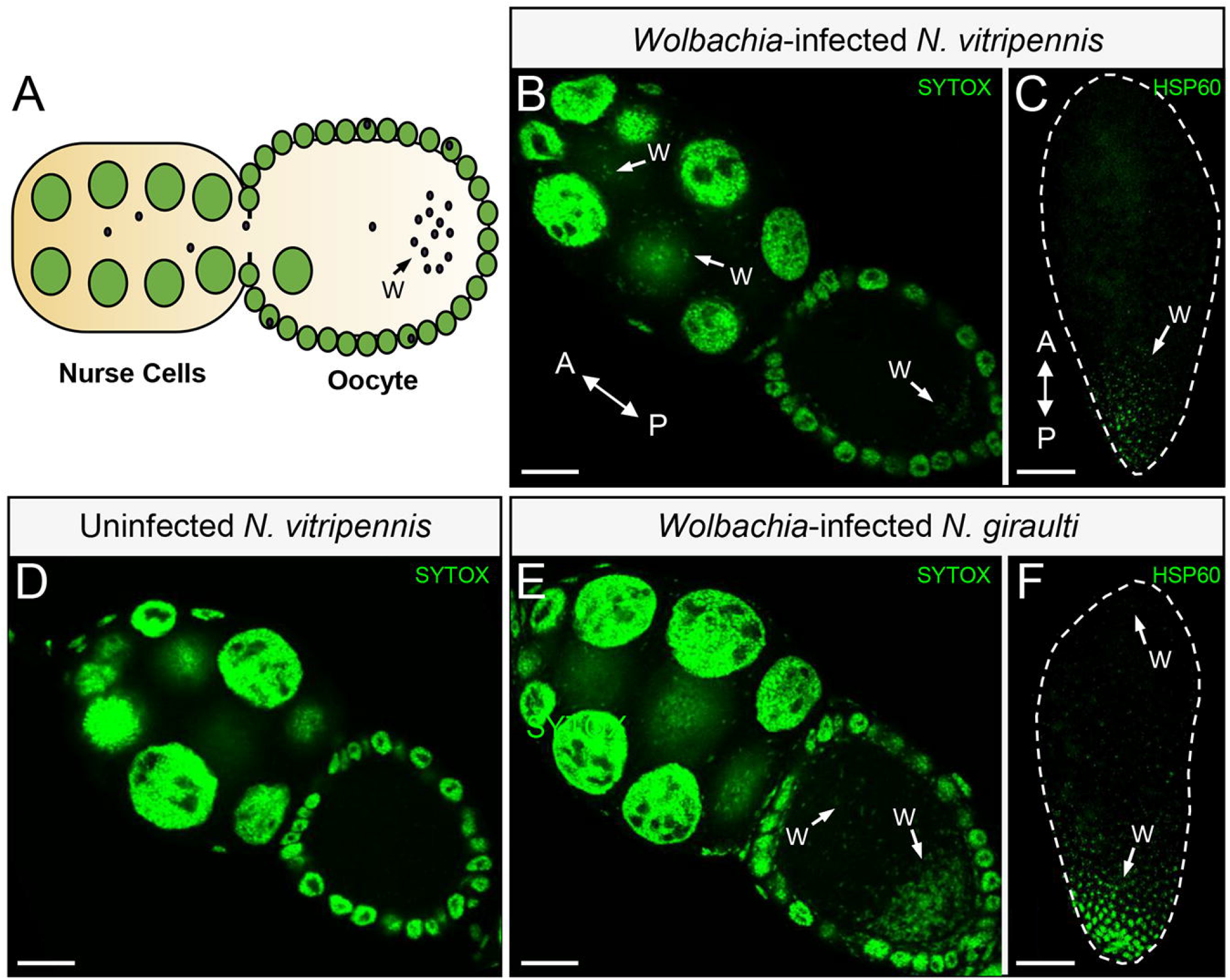
Disparities in *w*VitA titers begin during oogenesis. (A) Diagram of a *Nasonia* egg chamber. Large green circles represent nurse cell nuclei and small black circles represent *Wolbachia.* (B) Stage 3 egg chambers with host and *Wolbachia* DNA stained with SYTOX Green from *w*VitA-infected *N. vitripennis* 12.1. A = anterior, P = posterior, Scale bar = 15 µm. Examples of *Wolbachia* bacteria are labeled with a “W” and white arrows. (C) An embryo with host and *Wolbachia* DNA stained with HSP60 from *w*VitA-infected *N. vitripennis* 12.1. (D) Stage 3 egg chambers with host and *Wolbachia* DNA stained with SYTOX Green from uninfected *N. vitripennis* 12.1T. (E) Stage 3 egg chambers with host and *Wolbachia* DNA stained with SYTOX Green from *w*VitA-infected *N. giraulti* IntG. (F) An embryo with host and *Wolbachia* DNA stained with HSP60 from *w*VitA-infected *N. giraulti* IntG. All embryo and ovary images are representative of two and three independent experiments, respectively. See also Figure S1

### Phenotype-based selection and introgression identifies two maternal suppressor genomic regions

In an initial approach to determine the location and number of loci that suppress *w*VitA densities in *N. vitripennis*, we selected upon the low-density phenotype of this species while serially backcrossing hybrid females to *N. giraulti* IntG males (Figure 3A). Since the phenotype is controlled through a maternal genetic effect, hybrid females were selected based on the *w*VitA densities of their offspring, with sisters of the low-density offspring used as mothers in the next round of introgression (Figure 3A). Two independent selection lines were generated simultaneously to help discriminate between *N. vitripennis* regions maintained due to selection (present in both lines) versus those maintained through chance (present in only one line).

**Figure 3.**
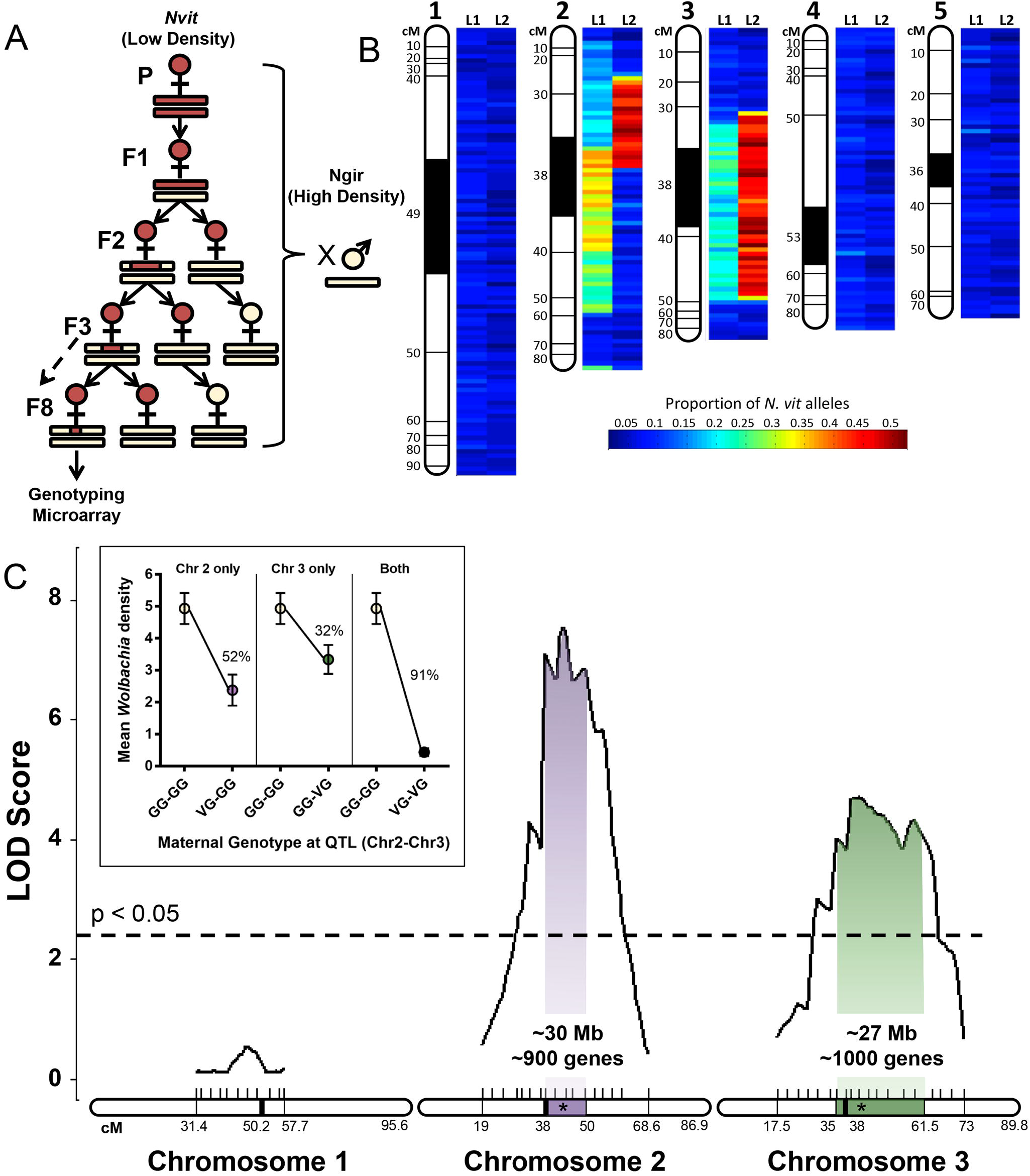
Two genomic regions interact additively to suppress *w*VitA titers. (A) Schematic of introgression using density phenotype (red female = low, cream female = high) as proxy for maternal genotype (red bar = *N. vitripennis*, cream bar = *N. giraulti*). Eighth generation females with lowest embryonic *Wolbachia* densities were genotyped on a *Nasonia* microarray. (B) Heatmap of the proportion of *N. vitripennis* alleles across the genome in a pool of three females from each introgression line (L1 or L2). The proportion of *N. vitripennis* alleles is scaled from 0 to 0.5, where 0 = no *N. vitripennis* alleles and 0.5 = all females were heterozygous. Areas were considered enriched for *N. vitripennis* alleles at ≥ 0.2. (C) Plot of LOD score after QTL mapping of F2 females. Shaded regions represent the 95% Bayes credible interval for significant QTL peaks (star). Dashed line represents genome-wide significance threshold at α = 0.05. (C, inset) *Wolbachia* density (mean ± S.E.M) of offspring based on maternal genotype at each QTL peak (V = *N. vitripennis*, G = *N. giraulti*). Percent reduction in densities is compared to offspring of F2 homozygous *N. giraulti* females. All maps are based on the *Nasonia* genetic map [40]. See also Table S1 and Data S1.

For each independent line, DNA from three females that produced ninth-generation offspring with the lowest *Wolbachia* densities were pooled and genotyped on a *Nasonia* genotyping microarray composed of 19,681 sequence markers that differ between *N. vitripennis* and *N. giraulti* [40]. Both selection lines (L1 and L2) displayed an enrichment of *N. vitripennis* alleles along the central portions of chromosomes 2 and 3 out of *Nasonia*’s five chromosomes (Figure 3B). On the most recent *N. vitripennis* linkage map [40], the area of enrichment on chromosome 2 for L1 occurs between 38 cM and 51.1 cM, while enrichment in L2 extends from 25.6 cM to 38 cM (Figure 3B). Although overlap in *N. vitripennis* allele enrichment between L1 and L2 on chromosome 2 occurs at 38 cM, the exact position and size of the overlap cannot be determined due to the fact that it falls within the poorly-assembled heterochromatic regions flanking the centromere [40]. For chromosome 3, the areas of enrichment for *N. vitripennis* alleles between L1 and L2 coincide starting at 35 cM and ending at 47.5 cM.

### QTL analysis validates the two maternal effect suppressor regions

To validate the *Wolbachia* density-suppressing chromosomal regions determined through phenotypic selection, we performed an independent quantitative trait loci (QTL) analysis in which F1 hybrid females were backcrossed to high-density *N. giraulti* IntG males to obtain 191 F2 recombinant females. Each F2 female was phenotyped by measuring the *Wolbachia* densities of her F3 pupal offspring. Since the most informative individuals in QTL mapping are those with the most extreme phenotypes [41], we selectively genotyped F2 females with the lowest (0.072 – 0.409, N = 42) and highest (2.958 – 10.674, N = 42) F3 pupal *Wolbachia* titers with a total of 47 microsatellite markers across chromosomes 1, 2 and 3 with an average distance between markers of 3 cM (Table S1). Using genotype data for selected individuals and phenotype data for all F2 females (Data S1), we identified two significant QTL regions at a genome-wide significance level of α = 0.05 (LOD > 2.29): one QTL peak on chromosome 2 at 43 cM (p < 0.001) and the other on chromosome 3 at 41.5 cM (p < 0.001, Figure 3C). Strikingly, the 95% Bayes credible interval on chromosome 2 corresponds to the same region identified by the genotyping microarray as enriched for *N. vitripennis* alleles in introgression Line 1 (38 cM – 51.1 cM), while the 95% Bayes credible interval on chromosome 3 also contains a region that was enriched for *N. vitripennis* alleles (35 cM – 47.5 cM) in both introgression lines. Thus, the microarray and QTL analyses complement each other and confirm that suppressor genes of major effect for *w*VitA density are located near the centromeric regions on chromosomes 2 and 3.

As a negative control, we genotyped the same individuals with markers located on *Nasonia* chromosome 1 (Data S1), which was not enriched for *N. vitripennis* alleles after the selection introgression. In the QTL analysis, the highest peak on chromosome 1 was not statistically significant (Figure 3C), indicating again that chromosomes 2 and 3 are likely the only chromosomes harboring genes of major effect for the *w*VitA density trait.

To determine the effect of each QTL on density suppression, the average percent reduction in F3 pupal *Wolbachia* densities was calculated for the F2 females with *N. vitripennis* alleles at markers close to one or both of the calculated QTL peaks. Females with *N. vitripennis* chromosome 2 or chromosome 3 QTLs produced offspring with a 52% or 32% reduction in densities, respectively, compared to offspring of females that were homozygous *N. giraulti* at both QTL loci (Figure 3C, inset). Furthermore, these effects acted additively for a 91% reduction in densities in offspring of females with *N. vitripennis* alleles at both loci compared to offspring of F2 females with *N. giraulti* alleles at both loci (Figure 3C, inset).

### Marker-assisted introgression confirms and narrows the maternal effect suppressor QTL on chromosome 3

To validate the QTLs on chromosomes 2 and 3 and narrow the gene candidate regions, we independently introgressed the QTL regions from *N. vitripennis* into an *N. giraulti* IntG background for at least nine generations using marker-assisted selection (similar to Figure 3A, see STAR Methods). After the ninth generation, we conducted sibling matings to produce segmental introgression lines that were homozygous *N. vitripennis* for the marker of interest. Unfortunately, generating *N. vitripennis* homozygous lines for the chromosome 2 region was not possible due to hybrid sterility, so we focused exclusively on the chromosome 3 region.

The initial homozygous and heterozygous introgression lines generated from sibling matings identified a candidate region 3.4 Mb in size containing 288 genes (Line IntC3) that suppressed *w*VitA densities by 60%, while lines lacking this region had little to no density suppression (Figure 4, Data S2). Surprisingly, the percent effect of the chromosome 3 homozygous introgression on *Wolbachia* suppression was nearly double that observed in the QTL study (60% vs 32%, Figure 3C inset). However, the QTL study was performed on F2 hybrid females while the introgression lines underwent at least nine generations of backcrossing. If there is a *N. vitripennis*-specific negative regulator of the *Wolbachia* suppressor gene on a different chromosome, then the allele would likely be present in F2 hybrids but would have recombined out with subsequent backcrossing to *N. giraulti* IntG. The stronger phenotype could also be due to the homozygous introgression lines having two copies of the *N. vitripennis* chromosome 3 candidate region, while F2 hybrid females were heterozygous. However, this is unlikely since heterozygous introgression females had the same level of *Wolbachia* suppression as their homozygous counterparts (Figure 4, C3-3 and C3-4 vs. C3-5 and C3-6, Kruskal-Wallis test: H = 1.39, df = 3, p = 0.71).

**Figure 4.**
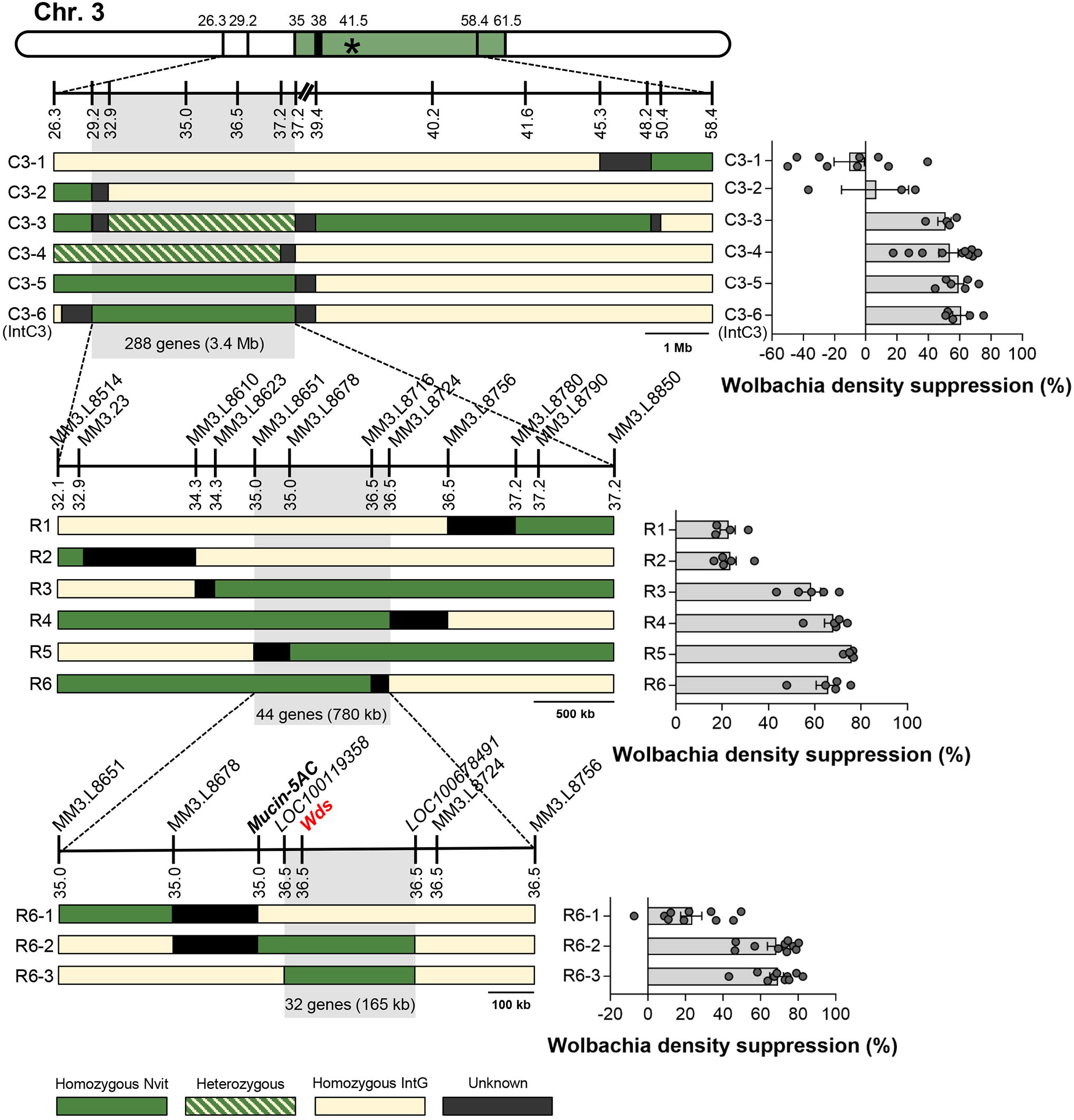
Segmental introgression lines narrow the chromosome 3 candidate region to 32 genes. Diploid genotypes are depicted as haplotypes, where green bars represent *N. vitripennis* homozygous regions, dashed bars are heterozygous regions, solid cream bars are *N. giraulti* IntG homozygous regions, and black bars are recombination breakpoints between two markers. The star and colored region on the chromosome map represent the QTL peak and 95% Bayes credible interval, respectively. Line graphs represent chromosome length in Mb and are drawn to scale except for centromeric regions (broken dashes at top of the figure). Names of the molecular markers (MM) used for genotyping are provided above the line graphs, and their locations in cM based on the genetic map from [40] are located below the line graphs. The bar graphs show the mean percent effect on density suppression in pupal offspring from all individual mothers with the same haplotype. Error bars denote mean ± S.E.M. See also Table S1, Table S3, and Data S2.

Line IntC3 was further backcrossed to *N. giraulti* IntG to generate four recombinant lines (R3, R4, R5, and R6) that suppressed *w*VitA densities by 58 – 78% (depending on the line) with an overlapping candidate region of 780 kb and 44 genes (Figure 4). Finally, line R6 was backcrossed to *N. giraulti* IntG to obtain three recombinant lines, two of which caused 67% (R6-2) and 68% (R6-3) density suppression. The overlapping *N. vitripennis* region in lines R6-2 and R6-3 was 165 kb and contained only 32 genes (Figure 4).

### RNA-seq identifies a single candidate gene (*Wds*) based on expression differences in ***Nasonia* ovaries**

To identify candidate genes within the 165 kb, 32-gene region that are differentially expressed in the maternal germline of *N. vitripennis* and *N. giraulti*, we performed high-throughput RNA sequencing (RNA-seq) on four independent pools of 40 ovary samples from the parental *N. vitripennis* line 12.1, the introgression line IntC3, and five independent pools from the parental *N. giraulti* line IntG (Table S2). Seven genes in the 32-gene candidate region exhibited significant differences in expression between the three aforementioned lines (Table S3). However, since the density trait is controlled by a dominant *N. vitripennis* maternal effect allele (Figure 1B), we reasoned that the most likely candidate gene(s) would be upregulated in *N. vitripennis* compared to *N. giraulti.* Analysis of the RNA-seq data indicated that only one of the seven genes (*LOC100679092*) was consistently and significantly overexpressed in *N. vitripennis* and IntC3 (low density) ovaries compared to *N. giraulti* IntG (high density) ovaries, which was confirmed in independent biological replicates by RT-qPCR (79-fold and 92-fold higher expression in *N. vitripennis* and IntC3 than *N. giraulti* IntG, respectively, Figure S2). The RNA-seq data also validated the same predicted gene splicing model for *LOC100679092* for both *N. vitripennis* and *N. giraulti*, indicating that the expression differences are not due to species-specific alternative splicing of the gene. As an uncharacterized gene with no known protein domains, we hereby name the gene *Wds* for *Wolbachia density suppressor* gene.

### *Wds* controls embryonic *w*VitA densities via a maternal effect

Parental RNAi has successfully been used in *Nasonia* to examine the effects of maternal genes on embryonic development [42, 43]. If the *N. vitripennis* allele of *Wds* (*Wds_v_*) is responsible for suppressing *Wolbachia* titers, we expect that post-transcriptional knockdown of *Wds_v_* transcripts in IntC3 mothers will result in reduced density suppression and, consequently, an increase in *w*VitA levels in the resulting embryos. Indeed, injection of IntC3 mothers with dsRNA against *Wds_v_*significantly increased offspring embryonic *w*VitA densities (696 ± 67.9, N = 24) by 56% or 63% compared to embryonic *w*VitA densities from mothers injected with dsRNA against a control bacterial gene, maltose transporter subunit E (*malE*) (447 ± 52.1, N = 24) or buffer-injected females (426 ± 50.3, N = 23), respectively (Figure 5A, Kruskal-Wallis test: H = 13.1, df = 3, p < 0.01, Dunn’s multiple comparisons test: p = 0.006 and p = 0.027 compared to *Wds_v_* group). This increase coincided with a 57% knock-down in *Wds_v_* gene expression in RNAi females compared to the buffer-injected controls (Figure 5B, Mann Whitney U test, p = 0.0015). Furthermore, we compared embryonic *w*VitA densities from mothers injected with dsRNA against *Nasonia* gene *LOC100679394* (*Mucin-5AC*), a gene that was significantly upregulated in *N. vitripennis* but immediately outside the chromosome 3 candidate region. Embryos from mothers injected with dsRNA against *Mucin-5AC* did not produce significantly higher *w*VitA densities (459 ± 75.9, N = 25) compared to embryos from either *malE*-RNAi (447 ± 52.1, N = 24) or buffer-injected females (426 ± 50.3, N = 25, Figure 5A), even though *Mucin-5AC-*RNAi mothers had a 71% decrease in *Mucin-5AC* gene expression versus buffer-injected controls (Figure S3, Mann Whitney U test p = 0.0003).

**Figure 5.**
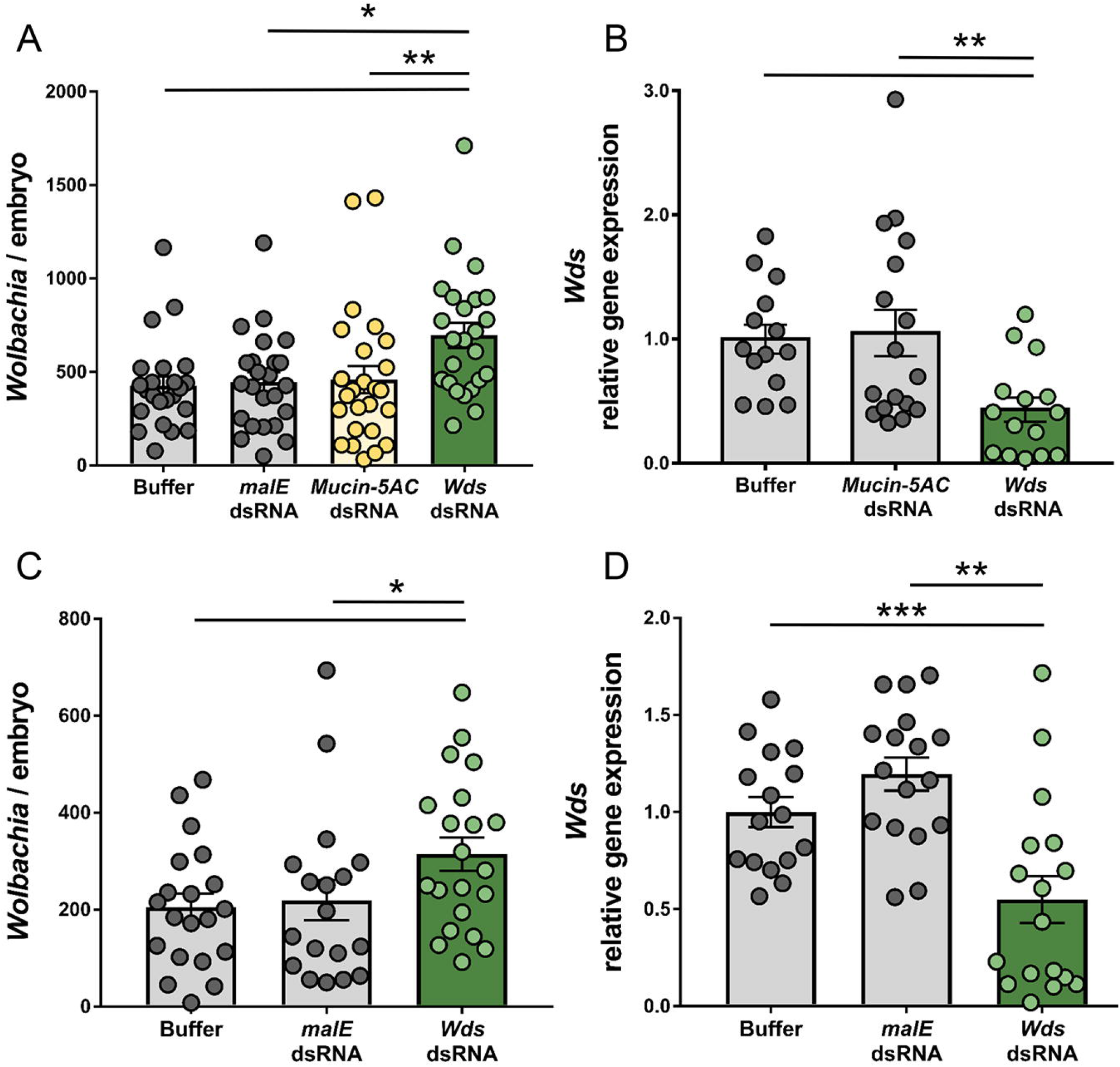
The *N. vitripennis* allele of *Wds* suppresses densities of vertically-transmitted *Wolbachia*. (A) Number of *w*VitA *Wolbachia* per embryo from IntC3 females that were buffer-injected, injected with dsRNA against control genes *MalE* or *Mucin-5AC*, or injected with dsRNA against *Wds_v_*. *p < 0.05 and **p < 0.01 post-hoc Dunn’s multiple comparisons test. (B) Relative gene expression of *Wds_v_* in late pupae of *Wds*-RNAi and *Mucin-5AC*-RNAi females normalized to *Wds_v_* expression in buffer-injected females. **p<0.01, Mann-Whitney U test. (C) Number of *w*VitA *Wolbachia* per embryo from R6-3 females that were buffer-injected, injected with dsRNA against control gene *MalE* or injected with dsRNA against *Wds_v_*. *p < 0.05, Mann-Whitney U test. (D) Relative gene expression of *Wds_v_*in late pupae of *Wds*-RNAi and *MalE*-RNAi females normalized to *Wds_v_* expression in buffer-injected females. **p<0.01, ***p < 0.001, Mann-Whitney U test. All error bars represent mean ± S.E.M. See also Figure S2, Figure S3, Table S2, Table S3, and Table S6.

To further validate the effect of *Wds_v_*on *Wolbachia* densities, females from recombinant line R6-3 (homozygous *N. vitripennis* for the 32-gene candidate region only) were injected with dsRNA against *Wds_v_.* Knock-down of *Wds_v_*in R6-3 females again significantly increased embryonic *w*VitA densities (314 ± 34.4, N = 21) by 43% or 54% compared to embryonic *w*VitA densities from mothers injected with dsRNA against the control bacterial gene *malE* (219 ± 39.2, N = 19) or from buffer-injected females (204 ± 28.4, N = 20), respectively (Figure 5C, Mann Whitney U, p = 0.049 and p = 0.023 compared to the *Wds_v_* group). This increase coincided with a 45% knock-down in *Wds_v_* gene expression in RNAi females compared to the buffer-injected controls (Figure 5D, Mann Whitney U test, p = 0.0041).

### Accelerated evolution and positive selection impacts *Wds* in *N. vitripennis*

The *Nasonia* genus is comprised of four closely-related species, with *N. vitripennis* sharing a common ancestor with the other three species approximately one million years ago [17, 18]. Wds protein sequences are 95% identical between *N. vitripennis* and *N. giraulti* with ten amino acid differences (and no indels) out of 201 total amino acids (Figure 6A). Between the more closely-related *N. giraulti* and *N. longicornis* species that diverged approximately 400,000 years ago [17], Wds is 99% identical with two amino acid differences that evolved specifically in *N. giraulti* (Figure 6A). Interestingly, Wds in *Trichomalopsis sarcophagae*, the wasp species most closely-related to *Nasonia* [44], shares 95% amino acid identity to the *N. vitripennis* protein, but 97% and 98% identity to the *N. giraulti* and *N. longicornis* proteins, respectively (Figure 6A). Taken together, there are seven unique amino acid changes in *N. vitripennis* that led to accelerated protein sequence evolution in *Wds_v_* (Figures 6A, B). Furthermore, three of those seven amino acid changes fall within a region of high positive selection based on a sliding window analysis of the Ka/Ks ratio (Figure 6C) [45]. Additionally, these changes are associated with a shift in the isoelectric point (pI) of the Wds protein, an important factor in protein evolution [46]. The pI drops from 9.24 in *N. vitripennis* to 8.75 in *N. giraulti*. In contrast, the isoelectric point difference for the Mucin-5AC control is minimal (ΔpI = 0.04).

**Figure 6.**
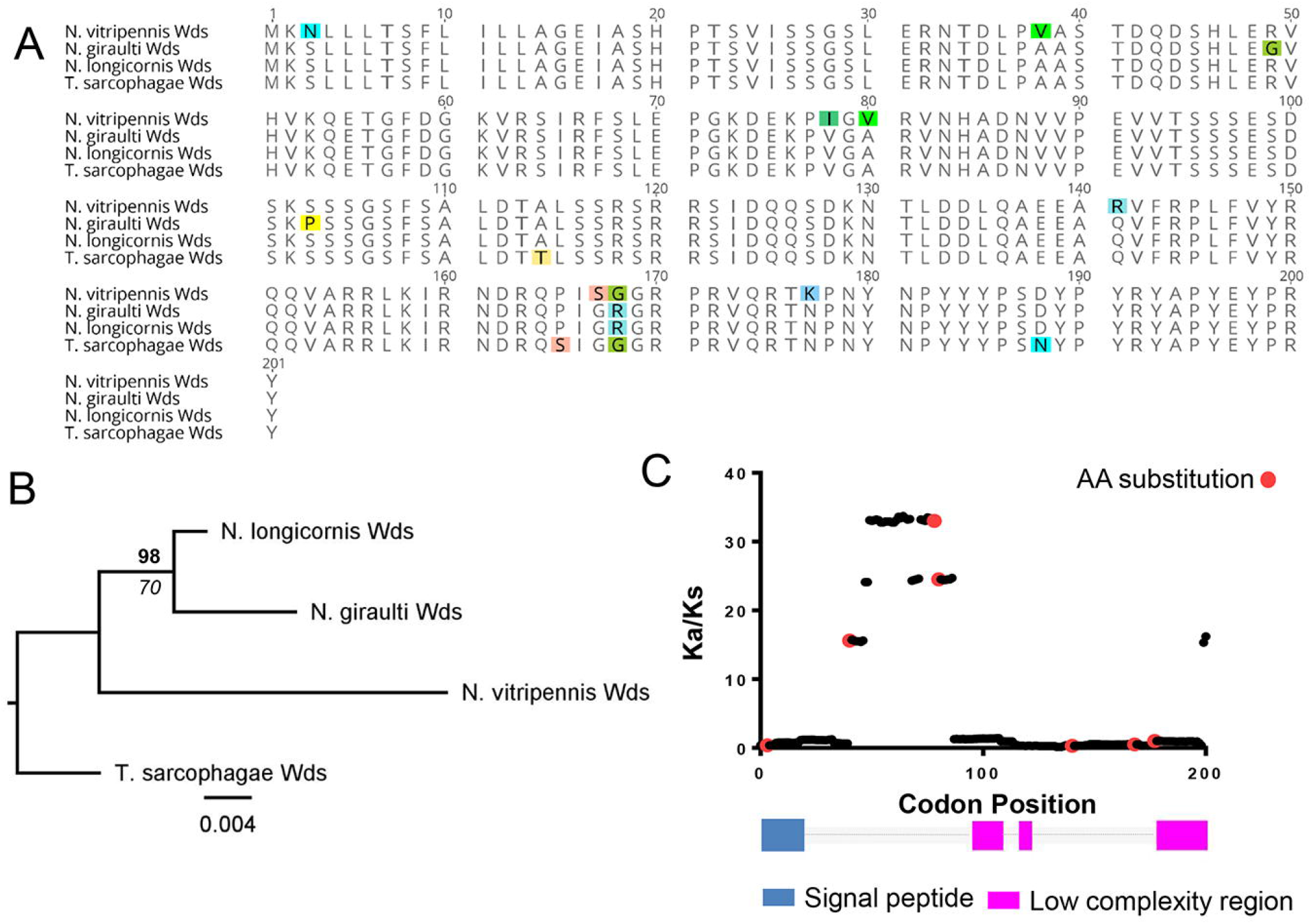
The *N. vitripennis* allele of *Wds* is under positive selection. (A) Alignment of Wds protein homologs from *N. vitripennis* (XP_008213336.1), *N. giraulti* (GL276173.1), *N. longicornis* (GL277955.1), and *Trichomalopsis sarcophagae* (OXU27029.1). (B) Amino acid phylogeny of Wds across the *Nasonia* genus and *Trichomalopsis sarcophagae*. Bold text above branch denotes Bayesian posterior probability. Italic text below branch denotes maximum likelihood bootstrap value. Scale bar denotes amino acid substitutions per site. (C) Plot of Ka/Ks ratios based on a sliding window analysis across the *N. vitripennis* and *N. giraulti Wds* coding sequences. Red circles indicate the locations of the seven amino acid substitutions unique to *N. vitripennis* Wds. Diagram below illustrates location of the predicted signal peptide and low complexity regions of the Wds protein. See also Table S4.

Overall, Wds in the *N. vitripennis* lineage experienced recent amino acid substitutions, possibly in response to acquisition of the *w*VitA *Wolbachia* strain that horizontally transferred into *N. vitripennis* after *N. vitripennis’* divergence from its common ancestor with *N. giraulti* and *N. longicornis* [39]. Outside of these four species, the next closest Wds orthologs are found in wasps such as *Trichogramma pretiosum, Copidosoma floridanum*, and *Polistes canadensis*, but they only share 29% – 42% amino acid identity to Wds_v_ across a majority of the sequence (Table S4). While more distant orthologs are present in other Hymenopterans such as bees and ants (Table S4), only portions of the proteins can be properly aligned. Therefore, our findings demonstrate a rapidly evolving, taxon-restricted gene can contribute directly to the adaptive evolution of regulating maternal symbiont transmission.

## DISCUSSION

The main goal of this study was to determine the number and types of gene(s) that control the most widespread, maternally-transmitted symbiont in animals [23, 24]. Unlike reverse genetic screens that mutate genes and then look for phenotypes, which may produce off-target effects unrelated to the true function of the protein, this forward genetic screen utilized an unbiased, candidate-blind approach to dissect the genetic basis of variation in host suppression of maternally-transmitted *Wolbachia.* We found that suppression of *w*VitA in *N. vitripennis* can be mapped to two regions of the *Nasonia* genome that regulate nearly all of the *Wolbachia* density suppression.

The identification of the *Wds* gene demonstrates that host regulation of maternally-transmitted symbiont density is adaptive and can proceed through lineage-specific amino acid changes in a maternal effect gene. The *N. vitripennis*-specific substitutions in Wds were possibly driven by *N. vitripennis’s* acquisition of *w*VitA after its divergence with *N. giraulti* [39]. Indeed, another *N. vitripennis*-specific *Wolbachia* strain, *w*VitB, maintains its low levels after transfer to *N. giraulti* [37], indicating that *Wds_v_*may encode a specific regulator of *w*VitA. Furthermore, *N. giraulti* maintains its native *w*GirA *Wolbachia* strain at comparable levels to *w*VitA in *N. vitripennis* [37], but whether *Wds_g_* is involved in regulating *w*GirA remains to be tested.

The Wds protein has areas of low complexity and a predicted signal peptide at its N-terminus (Figure 6C), but otherwise does not contain any characterized protein domains that allude to its function. Staining of *w*VitA in *Nasonia* ovaries revealed a trend of higher *Wolbachia* titers in the nurse cells of *N. vitripennis* than *N. giraulti* (Figure S1B,C) concurrent with significantly lower *Wolbachia* levels in *N. vitripennis* oocytes than in *N. giraulti* oocytes (Figure S1A). Thus, Wds_v_ may operate by hindering *w*VitA trafficking between nurse cells and the developing oocyte in *N. vitripennis*, perhaps by preventing *w*VitA binding to microtubule motor proteins responsible for *Wolbachia* transport into the oocyte [47, 48]. Furthermore, in *Drosophila, Wolbachia w*Mel bacteria in the oocyte increase proportionally faster than those in the nurse cells [47]. If the same is true for *Nasonia*, then high *w*VitA densities in *N. giraulti* could be a result of increased *w*VitA trafficking to the oocyte (due to a lack of repression by Wds_g_) compounded with faster proliferation once in the oocyte.

Alternatively, Wds_v_ could suppress *Wolbachia* replication by upregulating a host immune response or by downregulating host pathways that *Wolbachia* rely upon for growth. For example, inhibiting host proteasome activity in *Drosophila* significantly reduces *Wolbachia* oocyte titers [49], presumably due to a reduction in the availability of amino acids, a key nutrient that *Wolbachia* scavenges from its host [50]. However, if Wds_v_ regulates a general host pathway that impacts *Wolbachia* replication (such as host proteolysis), then we would expect both *w*VitA and *w*VitB titers to increase when transferred to *N. giraulti*. Instead, the strain-specificity of Wds_v_ suggests a more direct interaction with *Wolbachia*, such as a competitive inhibitor of motor protein binding and nurse cell to oocyte trafficking, as discussed above.

The findings presented here indicate that keeping maternally-transmitted symbionts in check can have a simple genetic basis, even for obligate intracellular bacteria that must be regulated within host cells and tissues. Moreover, a single maternal effect gene with a major consequence on the density phenotype demonstrates how natural selection can rapidly shape the evolution of density suppression of maternally transmitted symbionts in invertebrates. Future studies are warranted to tease apart the specific host-*Wolbachia* interactions driving *Wolbachia* regulation in *Nasonia* and to determine whether these interactions are paralleled in other insect-*Wolbachia* symbioses.

## ACKNOWLEDGMENTS

We would like to thank John Colbourne for his assistance with the *Nasonia* genotyping microarray, Jürgen Gadau for his helpful advice on QTL mapping, Jack Werren for providing the *Nasonia* genetic map before its publication, and Patrick Ferree for sharing his unpublished data of *Wolbachia* staining in *Nasonia* ovaries. We would also like to acknowledge Jeff Jones for helping with the MatLab heat-map figure. This work was supported by National Science Foundation award 1456778 to SRB and Graduate Research Fellowship 0909667 to LJF, National Institutes of Health awards (NIH) R01 AI132581 and R21 HD086833 to SRB, T32 GM008554, and a Vanderbilt Microbiome Initiative Award. Imaging was performed through the use of the Vanderbilt Cell Imaging Shared Resource (supported by NIH grants CA68485, DK20593, DK58404, DK59637 and EY08126). The funders had no role in study design, data collection and analysis, decision to publish, or preparation of the manuscript.

## AUTHOR CONTRIBUTIONS

Conceptualization, L.J.F., E.J.vO, and S.R.B.; Methodology, L.J.F., E.J.vO, and S.R.B.; Formal Analysis, L.J.F. and E.J.vO; Investigation, L.J.F., E.J.vO, and A.S.; Writing – Original Draft, L.J.F. and E.J.vO; Writing – Review and Editing, L.J.F., E.J.vO, A.S., and S.R.B.; Visualization, L.J.F. and E.J.vO. Project Administration, S.R.B.; Funding Acquisition, L.J.F. and S.R.B.

## STAR METHODS

### Contact for Reagent and Resource Sharing

Further information and requests for resources and reagents should be directed to and will be fulfilled by the Lead Contact, Seth R. Bordenstein.

### Experimental Model and Subject Details

#### *Nasonia* Parasitoid Wasps

Experiments were performed with *Nasonia vitripennis* strain 12.1, *N. giraulti* strain IntG12.1 or hybrids of these two species. *N. vitripennis* 12.1 is singly-infected with native *Wolbachia* strain *w*VitA and was derived from the double-infected *N. vitripennis* R511 (*w*VitA and *w*VitB) after a prolonged period of diapause [51]. *N. giraulti* strain IntG12.1 was generated by backcrossing *N. vitripennis* 12.1 females to uninfected *N. giraulti* Rv2x(u) males for nine generation [37], producing hybrids with an *N. giraulti* genome and an *N. vitripennis* cytoplasm harboring *w*VitA. All *Nasonia* were reared at 25°C in constant light on *Sarcophaga bullata* fly hosts reared in house on bovine liver from Walnut Hills Farm (Tennessee, USA).

### Method Details

#### Quantitative analysis of *Wolbachia* densities

Genomic DNA was extracted from pupae or adult *Nasonia* using the Gentra Puregene Tissue Kit (Qiagen) according to the manufacturer’s protocol. Real-time quantitative PCR (qPCR) was performed on a CFX96 Real-Time system (Bio-Rad) using a total reaction volume of 25 μl: 12.5 μl of iQ SYBR Green Supermix (Bio-Rad), 8.5 μl of sterile water, 1.0 μl each of 5 μM forward and reverse primers, and 2 μl of target DNA in single wells of a 96-well plate (Bio-Rad). All qPCR reactions were performed in technical duplicates and included a melt curve analysis to check for primer dimers and nonspecific amplification. Selective amplification was performed using primers previously described for the *Wolbachia groEL* gene [52] and *Nasonia NvS6K* gene [53]. Standard curves for each gene were constructed as previously described [53] using a log10 dilution series of larger PCR products of known concentrations for each gene. *groEL* and *S6K* copy numbers for each sample were calculated based on the following standard curve equations: *groEL*: y = −3.367x + 35.803 and *S6K:* y = −3.455x + 35.908, where y = averaged Ct value between technical duplicates and x = log starting quantity of template DNA. *Wolbachia* density was calculated by dividing *groEL* copy number by *S6K* copy number for each sample. Since diploid female *Nasonia* have twice the number of *S6K* copies than males, all experiments comparing *Wolbachia* densities were performed on either all male or all female samples to eliminate *S6K* copy number as a confounding factor in the statistical analyses.

#### Microsatellite marker genotyping

Primers used to amplify microsatellite markers that differ in size between *N. vitripennis* and *N. giraulti* are listed in Table S1. Microsatellite markers not previously published were identified by aligning *N. vitripennis* and *N. giraulti* genomic sequences using the Geneious alignment tool in Geneious Pro v5.5.8 (Biomatters). The Geneious primer design tool was then used to generate primer sets spanning each microsatellite. All PCR reactions were run on a Veriti Thermal Cycler (Applied Biosystems) with a total reaction volume of 15 μl: 7.5 μl of GoTaq Green Master Mix (Promega), 3.6 μl of sterile water, 1.2 μl of 5μM forward and reverse primers (see Table S1 for annealing temp.), and 1.5 μl of target DNA. PCR products were run on 4% agarose gels in TBE buffer (Sigma) at 90 volts for 2.5 to 6 hours, stained with GelRed (Biotium) according to manufacturer’s protocol, and imaged on a Red Personal Gel Imager (Alpha Innotech). New markers were validated based on predicted band size using *N. vitripennis* 12.1 and *N. giraulti* IntG as controls.

#### Phenotypic selection introgression and genotyping

*N. vitripennis* 12.1 females (low *w*VitA density) were backcrossed with *N. giraulti* IntG males (high *w*VitA density) for nine generations. For each generation of backcrossing, five female pupal offspring were pooled from each hybrid mother (N = 13 – 35 hybrid females depending on survival at each generation), and the pupal *Wolbachia* densities were measured using qPCR. Sisters of the pupae with the lowest *Wolbachia* densities were then used as mothers in the next round of backcrossing (N = 60 – 80 hybrid females). Two independent selection lines were maintained simultaneously along with control lines of pure-breeding *N. vitripennis* and *N. giraulti*. After eight generations of selection, the three females from each introgression line that produced ninth-generation offspring with the lowest *Wolbachia* densities were pooled and their DNA extracted using the DNeasy Blood and Tissue Kit (Qiagen) with the protocol for purification of DNA from insects. To obtain enough DNA for microarray hybridization, we used the REPLI-g Mini Kit (Qiagen) with the protocol for 5 μl of DNA template to amplify genomic DNA overnight at 30 °C, then purified the DNA using ethanol precipitation. The final concentration for each sample was diluted to 1 μg/μl and a total of 10 μl was sent to The Center for Genomics and Bioinformatics at Indiana University to be processed on a *Nasonia* genotyping microarray (Roche NimbleGen) tiled with probes for 19,681 single nucleotide polymorphisms and indels that differ between *N. vitripennis* and *N. giraulti* [40].

For each sample, the proportion of *N. vitripennis* alleles at each marker was determined based on the ratio of hybridization to the *N. vitripennis*-specific probe versus hybridization to the *N. giraulti*-specific probe, as previously described [40]. To verify species-specificity of these markers for our *Nasonia* strains, we also genotyped *N. vitripennis* 12.1 and *N. giraulti* IntG control females on the array, and markers that did not display the correct specificity within one standard deviation of the median were removed from subsequent analyses (5,301 markers total). The remaining markers were then manually mapped back to the most recent *Nasonia* linkage map [40]. Since all introgression females received one copy of their diploid genome from their *N. giraulti* father, the theoretical maximum proportion of *N. vitripennis* alleles at each marker cluster for experimental samples is 0.5. The proportion of *N. vitripennis* alleles was averaged for every 22 consecutive markers across each chromosome, and heat maps were generated using the HeatMap function in MATLAB (MathWorks). Areas were considered enriched for *N. vitripennis* alleles at ≥ 0.2.

#### QTL Anal**y**sis

F2 hybrid females (N = 191) were generated by backcrossing F1 *N. vitripennis*/*N. giraulti* hybrids to *N. giraulti* IntG males. F2 females were then backcrossed again to *N. giraulti* IntG and allowed to lay offspring. Five female pupae from each F2 female were pooled and their *Wolbachia* densities measured using qPCR. Females that produced offspring with densities within the highest and lowest quartile of the density distribution (N = 42 for each quartile) were selectively genotyped with 47 microsatellite markers spread across chromosomes 1, 2 and 3 with an average distance of 3 cM between markers (Table S1, Data S1). Phenotypic information for all 191 F2 females was included in the mapping analyses to prevent inflation of QTL effects due to the biased selection of extreme phenotypes [41]. QTL analyses were performed in R (version 3.0.2) with package R/qtl [54]. Significance thresholds for our dataset were calculated by using a stratified permutation test with the scanone function (1000 permutations). To identify significant QTL and their interactions, we first conducted a one-dimensional, one-QTL scan and a two-dimensional, two-QTL scan using the EM algorithm with a step size of 1 cM and an assumed genotype error probability of 0.001. Two significant QTLs were identified, one each on chromosomes 2 and 3, which were predicted to act additively. The positions of identified QTL were then refined using multiple QTL modeling with the multiple imputation algorithm (200 imputations, step size = 1 cM) assuming a model with two additive QTLs. 95% Bayes credible intervals were calculated for each QTL after multiple QTL modeling using the bayesint function.

#### Marker-assisted segmental introgressions

Marker-assisted segmental introgression lines were generated by repeatedly backcrossing hybrid females to *N. giraulti* IntG males for nine generations while selecting for *N. vitripennis* alleles at three microsatellite markers on chromosome 3 (MM3.17, NvC3-18, and MM3.37). After the ninth generation, families that maintained an *N. vitripennis* allele at one or more of these markers were selected, and siblings were mated to each other to produce lines containing homozygous *N. vitripennis* regions at and around the markers. Individual adult females from each segmental line were genotyped and phenotyped separately (N = 10 – 15 females per line). Females were hosted as virgins, five male pupal offspring per female were pooled, and pupal *Wolbachia* densities were measured using qPCR. Variation across plates for a single experiment was reduced by including a set of parental DNA controls on all plates. The parental fold-change was then calculated by dividing the average *N. giraulti* control density by the average *N. vitripennis* control density. To calculate the sample fold-change, the absolute density for each sample was divided by the average density of the *N. vitripennis* control. To determine how “effective” each segmental introgression line was at reducing densities, we calculated the percent effect on density suppression for each sample using the following equation:

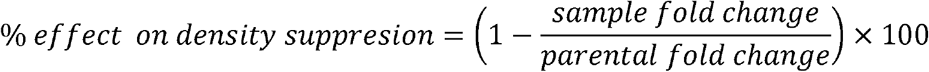

Each female was genotyped with markers across the region of interest, all females with identical genotypes across all markers were grouped together, and their percent effects on density suppression were averaged.

For the two subsequent rounds of introgression lines (R1 to R10 and R6-1 to R6-3), 300 IntC3 line females or 800 R6 line virgin females, respectively, were backcrossed to IntG males. The resulting virgin F1 females produced haploid, recombinant males. These recombinant males were mated to IntG females to produce heterozygous female offspring with the recombinant genotype. Siblings were then mated to each other and genotyped for two more generations to produce recombinant lines containing homozygous *N. vitripennis* introgressed regions reduced in size.

A subset of the genes within the 32-gene candidate region were genotyped using sequencing primers (Table S5) to amplify PCR products 250-750 bp for each of the R6 recombinant lines (R6-1 to R6-3) for Sanger sequencing (GENEWIZ). All PCR reactions were run on a Veriti Thermal Cycler (Applied Biosystems) with a total reaction volume of 15 μl: 7.5 μl of GoTaq Green Master Mix (Promega), 3.6 μl of sterile water, 1.2 μl of 5μM forward and reverse primers (see Table S5 for annealing temp.), and 1.5 μl of target DNA and purified using the QIAquick PCR Purification kit (Qiagen). At least 6 distinct single nucleotide polymorphisms (SNPs) between *N. vitripennis* and *N. giraulti* alleles were used to characterize the allele for each recombinant line using Geneious Pro v5.5.8 (Biomatters).

#### RNA-seq of ovaries

One-day old females from *Nasonia* strains *N. vitripennis* 12.1, *N. giraulti* IntG, and *N. vitripennis/N. giraulti* introgression line IntC3 were hosted as virgins on *S. bullata* pupae for 48 hours to stimulate feeding and oogenesis. Females were then dissected in RNase-free 1X PBS buffer, and their ovaries were immediately transferred to RNase-free Eppendorf tubes in liquid nitrogen. Forty ovaries were pooled for each replicate and 4-5 biological replicates were collected per *Nasonia* strain. Ovaries were manually homogenized with RNase-free pestles, and their RNA was extracted using the Nucleospin RNA/Protein Kit (Macherey-Nagel) according to the manufacturer’s protocol for purification of total RNA from animal tissues. After RNA purification, samples were treated with RQ1 RNase-free DNase (Promega) for 1 hour at 37 °C, followed by an ethanol precipitation with 1/10^th^ volume 3M sodium acetate and 3 volumes 100% ethanol incubated overnight at −20 °C. PCR of samples with *Nasonia* primers NvS6KQTF4 and NVS6KQTR4 [53] revealed some residual DNA contamination, so DNase treatment and ethanol precipitation were repeated. After the second DNase treatment, PCR with the same primer set confirmed absence of contaminating DNA. Sample RNA concentrations were measured with a Qubit 2.0 Fluorometer (Life Technologies) using the RNA HS Assay kit (Life Technologies). All samples were run multiplexed on two lanes of the Illumina HiSeq3000 (paired-end, 150 bp reads, ∼30M reads) at Vanderbilt’s VANTAGE sequencing core. Raw reads were trimmed and mapped to the *N. vitripennis* genome Nvit_2.1 (GCF_000002325.3) in CLC Genomics Workbench 8.5.1, allowing ten gene hits per read using a minimum length fraction of 0.9 and a minimum similarity fraction of 0.9. The number of reads generated for each sample and the percentage of reads that mapped to the *N. vitripennis* genic and intergenic regions are provided in Table S2. Significant differential gene expression was determined in CLC Genomics Workbench 8.5.1 at α = 0.05 for unique gene reads using the Empirical analysis of DGE tool, which is based on the edgeR program commonly used for gene expression analyses [55].

#### RT-qPCR validation of RNA-seq results

One-day old females from *N. vitripennis* 12.1, *N. giraulti* IntG, and IntC3 were hosted with two *S. bullata* pupae and honey to encourage ovary development. After 48 hours, ovaries were removed in RNase-free PBS, flash-frozen in liquid nitrogen then stored at −80 °C. 4-5 replicates of fifty ovaries per replicate were collected for each *Nasonia* strain. Total RNA was extracted from each sample using Trizol reagent (Invitrogen) with the Direct-zol RNA Miniprep kit (Zymo Research) then treated with the DNA-free DNA Removal kit (Ambion) for one hour at 37 °C. After ensuring with PCR that all DNA had been removed, RNA was converted to cDNA using the SuperScript VILO cDNA Synthesis kit (Invitrogen).

RT-qPCR was performed on a CFX96 Real-Time system (Bio-Rad) using a total reaction volume of 25 μl: 12.5 μl of iTaq Universal SYBR Green Supermix (Bio-Rad), 8.5 μl of sterile water, 1 μl each of 5 μM forward and reverse primers (see STAR Methods), and 2 μl of target cDNA in single wells of a 96-well plate (Bio-Rad). All RT-qPCR reactions were performed in technical duplicates and included a melt curve analysis to check for nonspecific amplification. The 60S ribosomal protein L32 (also known as RP49) was used as an expression control. All primers for RT-qPCR are provided in Table S6. Expression values for each candidate gene were calculated using the ΔΔC_t_ method of relative quantification [56] with RP49 as the reference gene. Fold-change was determined by normalizing expression values to the mean expression value of *N. giraulti* IntG for each gene.

#### RNAi of candidate genes

To generate DNA template for dsRNA synthesis, primers with a T7 promoter sequence on the 5’ end of each primer were used to amplify a 450-700 bp region of the targeted genes (Table S6) by PCR using *N. vitripennis* whole-body cDNA as template. PCR amplicons were separated by electrophoresis on a 1% agarose gel, excised, and purified using the QIAquick Gel Extraction kit (Qiagen). The purified PCR products were used as template for a second PCR reaction with the same gene-specific T7 primers, then purified using the QIAquick PCR Purification kit (Qiagen). After quantification with the Qubit dsDNA Broad Range Assay kit (Thermo Fisher Scientific), approximately 1 ug of the purified PCR amplicon was used as template for dsRNA synthesis with the MEGAScript RNAi kit (Ambion). The dsRNA synthesis reaction was incubated for six hours at 37 °C, treated with RNase and DNase for one hour at 37 °C, then column-purified according to the manufacturer’s protocol.

For injection, the dsRNA was used at a final concentration standardized to 750 ng/ul dsRNA. A Nanoject II (Drummond Scientific) was used to inject 23 nl of dsRNA (or MEGAScript kit elution buffer) into the ventral abdomen of female *Nasonia* at the yellow pupal stage. After emerging as adults, injected females were given honey and hosted individually on two *S. bullata* pupae for 48 hours. On the third day after emergence, they were transferred to new vials where they were presented with a single *S. bullata* host. After five hours, the hosts were opened and up to ten embryos were collected in a 1.5 ml Eppendorf tube for each female and stored at −80 °C. The females were given two hosts overnight, and then the same process was repeated again on the fourth day.

The number of *Wolbachia* bacteria per embryo from injected females three and four days post emergence was determined using qPCR with *Wolbachia groEL* primers as described above. *Wolbachia* titers were not normalized to *Nasonia* gene copy number because early embryos have varying numbers of genome copies depending on how many rounds of mitotic division they have undergone [57]. To determine the knock-down efficiency of each dsRNA injection, RNA extraction and RT-qPCR of black pupae, five days post injection, were performed with 14-17 biological replicates from each treatment group as described above using the gene-specific RT-qPCR primers in Table S6.

#### *Wds* phylogeny and selection analyses

*N. giraulti* and *N. longicornis* nucleotide sequences of *Wds* were obtained from NCBI genomic scaffold sequences GL276173 and GL277955, respectively, and indels were manually extracted in Geneious Pro v11.0.3 (Biomatters) based on homology to *N. vitripennis* gene LOC1006079092. Protein alignment of Wds amino acid sequences for the three *Nasonia spp.* and its homolog in *T. sarcophagae* (TSAR_005991) was performed using the Geneious alignment tool. MEGA7.0.26 [58] was used to identify the JTT model as the best model of protein evolution for the alignment based on corrected Akaike information criterion (AICc). PhyML [59] and MrBayes [60] were executed in Geneious with default parameters to construct a maximum likelihood tree with bootstrapping and a Bayesian tree with a burn-in of 100,000, respectively.

To identify residues under positive selection in Wds, Ka/Ks values were calculated based on a pairwise alignment of the *N. vitripennis* and *N. giraulti Wds* coding sequences using a sliding window analysis (window = 30 AA, step size = 1 AA, Standard Code for genetic code input) in the SWAKK bioinformatics web server [45]. Analysis of Wds for protein structures and conserved domains was performed using the SMART online software at http://smart.embl-heidelberg.de [61]. Protein pIs were predicted using the online ExPASY Compute pI/Mw tool.

#### Imaging *Wolbachia* in *Nasonia*

For *Wolbachia* staining in ovaries, female *Nasonia* were hosted on *Sarcophaga bullata* pupae for two to three days before dissection to encourage ovary development. Females were dissected in 1X phosphate-buffered saline (PBS) solution, where ovaries were removed with forceps and individual ovarioles were separated with fine needles. Ovaries were fixed in 4% formaldehyde in PBS with 0.2% Triton X-100 (PBST) for 20 minutes at room temperature then transferred to a 1.5 ml Eppendorf tube containing PBST. Samples were washed quickly three times with PBST then incubated in PBST plus 1 mg/ml RNase A for three hours at room temperature then overnight at 4 °C. After removing the RNase A solution, ovaries were incubated at room temperature for 15 minutes in PBST with 1:300 SYTOX green nucleic acid stain (Thermo Fisher Scientific) before washing twice with PBST, 15 minutes each time. Ovaries were then transferred to a glass slide and mounted in ProLong Gold antifade solution (Thermo Fisher Scientific) and covered with a glass cover slip sealed with nail polish. Ovary images in Figure 2 are representative of three independent experiments performed on different days with 2 – 3 females per species.

For *Wolbachia* staining in embryos, female *Nasonia* were hosted on a single *S. bullata* pupae for five hours. The host puparium was peeled away, and embryos were transferred with a probe to a glass vial containing 5 mL heptane. After shaking for two minutes, 5 mL methanol was added to the vial and shaken for another two minutes. Dechorionated embryos that sunk to the bottom of the vial were transferred to a 1.5 mL Eppendorf tube with methanol then were serially rehydrated in increasing ratios of methanol to PBST for 1 min each (90% MeOH:10% PBST, 75% MeOH: 25% PBST, 50% MeOH: 50% PBST, 25% MeOH:75% PBST) before a final wash in 100% PBST for 5 mins. Embryos were then blocked in PBST + 0.2% BSA (PBST-BSA) for 30 mins then PBST-BSA + 5% normal goat serum (PBANG) for 1 hour followed by a 2-hour incubation in PBANG + 1 mg/ml RNase. Embryos were stained overnight at 4 °C with monoclonal mouse anti-human Hsp60 antibody (Sigma; 1:250), which cross-reacts with *Wolbachia* but not insect proteins [62, 63]. After washing in PBST-BSA (4X, 15 mins each), embryos were incubated in goat anti-mouse Alexa Fluor 594 (Thermo Fisher Scientific; 1:500) for 2 hours, washed again in PBST-BSA (4X, 15 mins each), then stained with SYTOX Green (1:300) nuclear stain. Embryos were mounted to coverslips with Prolong Diamond Antifade Mountant (Thermo Fisher Scientific). Embryo images in Figure 2 are representative of two independent experiments performed on different days.

All images were acquired on a Zeiss LSM 510 META inverted confocal microscope at the Vanderbilt University Medical Center Cell Imaging Shared Resource core and processed with Fiji software [64]. Quantification of *Wolbachia* in *Nasonia* oocytes and nurse cells of stage 3 egg chambers was performed by calculating corrected total cell fluorescence with ImageJ software 1.47v. *Nasonia* oocyte images taken at the same relative z-stack slice were traced and the area integrated intensity was measured and compared to a background region (a traced region next to the egg chamber that has no fluorescence). Corrected total cell fluorescence was calculated as integrated intensity of the oocyte – (area of the oocyte * mean fluorescence of the background region). Quantification of *Wolbachia* in *Nasonia* nurse cells was also measured by counting the total number of fluorescent puncta, representative of *Wolbachia* and absent in *Wolbachia*-free *Nasonia*, in the *Nasonia* nurse cell cytoplasms.

### Quantification and Statistical Analyses

All statistical analyses, unless otherwise noted, were performed in GraphPad Prism 6.07 (GraphPad Software, La Jolla, CA). Outliers were removed from the results on embryonic *Wolbachia* titers and *Wds* RT-qPCR using the ROUT method, Q=1%. Non-parametric tests were used on all data since most data did not pass a Shapiro-Wilk test for normality or sample sizes were too small. Mann-Whitney U tests were used for comparisons between two groups, whereas a Kruskal-Wallis test was used to compare multiple groups. If the Kruskal-Wallis test was significant (p ≤ 0.05), a post-hoc Dunn’s test of multiple comparisons was used to calculate significance for all pair-wise combinations within the group. All averages are reported as mean ± S.E.M. For all quantifications of pupal *Wolbachia* densities, sample size “N” represents one pool of five pupae. For any data referring to adult *Nasonia* (genotyping, RNAi or RT-qPCR), sample size “N” denotes individual *Nasonia*. Statistical parameters for each experiment are reported in the results section and in any related figure legends.

### Data and Software Availability

RNA-seq data generated in this study is available in the Sequence Reads Archive under Bioproject ID PRJNA430433.

### SUPPLEMENTAL INFORMATION

Supplemental Information includes three figures, six tables and two datasets.

**Data S1. QTL genotypes and phenotypes, Related to Figure 3.**

V/G = heterozygous female (red), G/G = homozygous *N. giraulti* female (cream), and question marks indicate an unknown genotype. Markers that fall within the 95% Bayes cedible interval are highlighted in purple (chromosome two) or green (chromosome three).

**Data S2. Chromosome 3 segmental introgression haplotypes and their effects on *w*VitA density suppression, Related to Figure 4.**

cM locations based on genetic linkage map from [40].

## REFERENCES

1. Ley, R.E., Turnbaugh, P.J., Klein, S., and Gordon, J.I. (2006). Microbial ecology: human gut microbes associated with obesity. Nature 444, 1022–1023.

2. Turnbaugh, P.J., Ley, R.E., Mahowald, M.A., Magrini, V., Mardis, E.R., and Gordon, J.I. (2006). An obesity-associated gut microbiome with increased capacity for energy harvest. Nature 444, 1027–1031.

3. Round, J.L., Lee, S.M., Li, J., Tran, G., Jabri, B., Chatila, T.A., and Mazmanian, S.K. (2011). The Toll-like receptor 2 pathway establishes colonization by a commensal of the human microbiota. Science 332, 974–977.

4. Ivanov, II, Frutos Rde, L., Manel, N., Yoshinaga, K., Rifkin, D.B., Sartor, R.B., Finlay, B.B., and Littman, D.R. (2008). Specific microbiota direct the differentiation of IL-17-producing T-helper cells in the mucosa of the small intestine. Cell Host Microbe 4, 337–349.

5. Candela, M., Perna, F., Carnevali, P., Vitali, B., Ciati, R., Gionchetti, P., Rizzello, F., Campieri, M., and Brigidi, P. (2008). Interaction of probiotic *Lactobacillus* and *Bifidobacterium* strains with human intestinal epithelial cells: adhesion properties, competition against enteropathogens and modulation of IL-8 production. Int J Food Microbiol 125, 286–292.

6. Fukuda, S., Toh, H., Hase, K., Oshima, K., Nakanishi, Y., Yoshimura, K., Tobe, T., Clarke, J.M., Topping, D.L., Suzuki, T., et al. (2011). Bifidobacteria can protect from enteropathogenic infection through production of acetate. Nature 469, 543–547.

7. Calderone, R.A., and Fonzi, W.A. (2001). Virulence factors of *Candida albicans*. Trends Microbiol 9, 327–335.

8. Mitchell, J. (2011). *Streptococcus mitis*: walking the line between commensalism and pathogenesis. Mol Oral Microbiol 26, 89–98.

9. Fleury, F., Vavre, F., Ris, N., Fouillet, P., and Bouletreau, M. (2000). Physiological cost induced by the maternally-transmitted endosymbiont *Wolbachia* in the *Drosophila* parasitoid *Leptopilina heterotoma*. Parasitology 121 *Pt* *5*, 493–500.

10. Min, K.T., and Benzer, S. (1997). *Wolbachia*, normally a symbiont of *Drosophila*, can be virulent, causing degeneration and early death. Proc Natl Acad Sci U S A 94, 10792–10796.

11. Koehncke, A., Telschow, A., Werren, J.H., and Hammerstein, P. (2009). Life and death of an influential passenger: *Wolbachia* and the evolution of CI-modifiers by their hosts. PLoS One 4, e4425.

12. Jaenike, J. (2009). Coupled population dynamics of endosymbionts within and between hosts. Oikos 118, 353–362.

13. Funkhouser, L.J., and Bordenstein, S.R. (2013). Mom knows best: the universality of maternal microbial transmission. PLoS Biol 11, e1001631.

14. Login, F.H., Balmand, S., Vallier, A., Vincent-Monegat, C., Vigneron, A., Weiss-Gayet, M., Rochat, D., and Heddi, A. (2011). Antimicrobial peptides keep insect endosymbionts under control. Science 334, 362–365.

15. Serbus, L.R., Ferreccio, A., Zhukova, M., McMorris, C.L., Kiseleva, E., and Sullivan, W. (2011). A feedback loop between *Wolbachia* and the *Drosophila* gurken mRNP complex influences *Wolbachia* titer. J Cell Sci 124, 4299–4308.

16. Newton, I.L., Savytskyy, O., and Sheehan, K.B. (2015). *Wolbachia* utilize host actin for efficient maternal transmission in *Drosophila melanogaster*. PLoS Pathog 11, e1004798.

17. Campbell, B.C., Steffen-Campbell, J.D., and Werren, J.H. (1993). Phylogeny of the *Nasonia* species complex (Hymenoptera: Pteromalidae) inferred from an internal transcribed spacer (ITS2) and 28S rDNA sequences. Insect Mol Biol 2, 225–237.

18. Raychoudhury, R., Desjardins, C.A., Buellesbach, J., Loehlin, D.W., Grillenberger, B.K., Beukeboom, L., Schmitt, T., and Werren, J.H. (2010). Behavioral and genetic characteristics of a new species of *Nasonia*. Heredity 104, 278–288.

19. Gadau, J., Page, R.E., and Werren, J.H. (2002). The genetic basis of the interspecific differences in wing size in *Nasonia* (Hymenoptera; Pteromalidae): major quantitative trait loci and epistasis. Genetics 161, 673–684.

20. Loehlin, D.W., Oliveira, D.C., Edwards, R., Giebel, J.D., Clark, M.E., Cattani, M.V., van de Zande, L., Verhulst, E.C., Beukeboom, L.W., Munoz-Torres, M., et al. (2010). Non-coding changes cause sex-specific wing size differences between closely related species of *Nasonia*. PLoS Genet 6, e1000821.

21. Werren, J.H., Cohen, L.B., Gadau, J., Ponce, R., Baudry, E., and Lynch, J.A. (2015). Dissection of the complex genetic basis of craniofacial anomalies using haploid genetics and interspecies hybrids in *Nasonia* wasps. Dev Biol 415, 391–405.

22. Niehuis, O., Buellesbach, J., Gibson, J.D., Pothmann, D., Hanner, C., Mutti, N.S., Judson, A.K., Gadau, J., Ruther, J., and Schmitt, T. (2013). Behavioural and genetic analyses of *Nasonia* shed light on the evolution of sex pheromones. Nature 494, 345–348.

23. Zug, R., and Hammerstein, P. (2012). Still a host of hosts for *Wolbachia*: analysis of recent data suggests that 40% of terrestrial arthropod species are infected. PLoS One 7, e38544.

24. Weinert, L.A., Araujo-Jnr, E.V., Ahmed, M.Z., and Welch, J.J. (2015). The incidence of bacterial endosymbionts in terrestrial arthropods. Proc Biol Sci 282, 20150249.

25. Werren, J.H., Zhang, W., and Guo, L.R. (1995). Evolution and Phylogeny of *Wolbachia*-Reproductive Parasites of Arthropods. Proc R Soc Lond B Biol Sci 261, 55–63.

26. Vavre, F., Fleury, F., Lepetit, D., Fouillet, P., and Bouletreau, M. (1999). Phylogenetic evidence for horizontal transmission of *Wolbachia* in host-parasitoid associations. Mol Biol Evol 16, 1711–1723.

27. Werren, J.H., Baldo, L., and Clark, M.E. (2008). *Wolbachia*: master manipulators of invertebrate biology. Nat Rev Microbiol 6, 741–751.

28. Serbus, L.R., Casper-Lindley, C., Landmann, F., and Sullivan, W. (2008). The genetics and cell biology of *Wolbachia*-host interactions. Annu Rev Genet 42, 683–707.

29. Perrot-Minnot, M.J., and Werren, J.H. (1999). *Wolbachia* infection and incompatibility dynamics in experimental selection lines. J Evol Biol 12, 272–282.

30. Dyer, K.A., Minhas, M.S., and Jaenike, J. (2005). Expression and modulation of embryonic male-killing in *Drosophila innubila*: opportunities for multilevel selection. Evolution 59, 838–848.

31. McMeniman, C.J., Lane, R.V., Cass, B.N., Fong, A.W., Sidhu, M., Wang, Y.F., and O’Neill, S.L. (2009). Stable introduction of a life-shortening *Wolbachia* infection into the mosquito *Aedes aegypti*. Science 323, 141–144.

32. Suh, E., Mercer, D.R., Fu, Y., and Dobson, S.L. (2009). Pathogenicity of life-shortening *Wolbachia* in *Aedes albopictus* after transfer from *Drosophila melanogaster*. Appl Environ Microbiol 75, 7783–7788.

33. Le Clec’h, W., Braquart-Varnier, C., Raimond, M., Ferdy, J.B., Bouchon, D., and Sicard, M. (2012). High virulence of *Wolbachia* after host switching: when autophagy hurts. PLoS Pathog 8, e1002844.

34. Mouton, L., Henri, H., Bouletreau, M., and Vavre, F. (2003). Strain-specific regulation of intracellular *Wolbachia* density in multiply infected insects. Mol Ecol 12, 3459–3465.

35. Kondo, N., Shimada, M., and Fukatsu, T. (2005). Infection density of *Wolbachia* endosymbiont affected by co-infection and host genotype. Biol Lett 1, 488–491.

36. Rio, R.V., Wu, Y.N., Filardo, G., and Aksoy, S. (2006). Dynamics of multiple symbiont density regulation during host development: tsetse fly and its microbial flora. Proc Biol Sci 273, 805–814.

37. Chafee, M.E., Zecher, C.N., Gourley, M.L., Schmidt, V.T., Chen, J.H., Bordenstein, S.R., and Clark, M.E. (2011). Decoupling of host-symbiont-phage coadaptations following transfer between insect species. Genetics 187, 203–215.

38. Bian, G., Zhou, G., Lu, P., and Xi, Z. (2013). Replacing a native *Wolbachia* with a novel strain results in an increase in endosymbiont load and resistance to dengue virus in a mosquito vector. PLoS Negl Trop Dis 7, e2250.

39. Raychoudhury, R., Baldo, L., Oliveira, D.C., and Werren, J.H. (2009). Modes of acquisition of *Wolbachia:* horizontal transfer, hybrid introgression, and codivergence in the *Nasonia* species complex. Evolution 63, 165–183.

40. Desjardins, C.A., Gadau, J., Lopez, J.A., Niehuis, O., Avery, A.R., Loehlin, D.W., Richards, S., Colbourne, J.K., and Werren, J.H. (2013). Fine-scale mapping of the *Nasonia* genome to chromosomes using a high-density genotyping microarray. G3 *3*, 205–215.

41. Lander, E.S., and Botstein, D. (1989). Mapping mendelian factors underlying quantitative traits using RFLP linkage maps. Genetics 121, 185–199.

42. Lynch, J.A., Brent, A.E., Leaf, D.S., Pultz, M.A., and Desplan, C. (2006). Localized maternal orthodenticle patterns anterior and posterior in the long germ wasp *Nasonia*. Nature 439, 728–732.

43. Verhulst, E.C., Beukeboom, L.W., and van de Zande, L. (2010). Maternal control of haplodiploid sex determination in the wasp *Nasonia*. Science 328, 620–623.

44. Werren, J.H., and Loehlin, D.W. (2009). The parasitoid wasp *Nasonia*: an emerging model system with haploid male genetics. Cold Spring Harb Protoc 2009, pdb emo134.

45. Liang, H., Zhou, W., and Landweber, L.F. (2006). SWAKK: a web server for detecting positive selection in proteins using a sliding window substitution rate analysis. Nucleic Acids Res 34, W382–384.

46. Alende, N., Nielsen, J.E., Shields, D.C., and Khaldi, N. (2011). Evolution of the isoelectric point of mammalian proteins as a consequence of indels and adaptive evolution. Proteins 79, 1635–1648.

47. Ferree, P.M., Frydman, H.M., Li, J.M., Cao, J., Wieschaus, E., and Sullivan, W. (2005). *Wolbachia* utilizes host microtubules and Dynein for anterior localization in the *Drosophila* oocyte. PLoS Pathog 1, e14.

48. Serbus, L.R., and Sullivan, W. (2007). A cellular basis for *Wolbachia* recruitment to the host germline. PLoS Pathog 3, e190.

49. White, P.M., Serbus, L.R., Debec, A., Codina, A., Bray, W., Guichet, A., Lokey, R.S., and Sullivan, W. (2017). Reliance of *Wolbachia* on High Rates of Host Proteolysis Revealed by a Genome-Wide RNAi Screen of Drosophila Cells. Genetics 205, 1473–1488.

50. Caragata, E.P., Rances, E., O’Neill, S.L., and McGraw, E.A. (2014). Competition for amino acids between *Wolbachia* and the mosquito host, *Aedes aegypti*. Microb Ecol 67, 205–218.

51. Perrot-Minnot, M.J., Guo, L.R., and Werren, J.H. (1996). Single and double infections with Wolbachia in the parasitic wasp Nasonia vitripennis: effects on compatibility. Genetics 143, 961–972.

52. Bordenstein, S.R., Marshall, M.L., Fry, A.J., Kim, U., and Wernegreen, J.J. (2006). The tripartite associations between bacteriophage, *Wolbachia*, and arthropods. PLoS Pathog 2, e43.

53. Bordenstein, S.R., and Bordenstein, S.R. (2011). Temperature affects the tripartite interactions between bacteriophage WO, *Wolbachia*, and cytoplasmic incompatibility. PLoS One 6, e29106.

54. Broman, K.W., Wu, H., Sen, S., and Churchill, G.A. (2003). R/qtl: QTL mapping in experimental crosses. Bioinformatics 19, 889–890.

55. Robinson, M.D., McCarthy, D.J., and Smyth, G.K. (2010). edgeR: a Bioconductor package for differential expression analysis of digital gene expression data. Bioinformatics 26, 139–140.

56. Livak, K.J., and Schmittgen, T.D. (2001). Analysis of relative gene expression data using real-time quantitative PCR and the 2(-Delta Delta C(T)) Method. Methods 25, 402–408.

57. Pultz, M.A., Westendorf, L., Gale, S.D., Hawkins, K., Lynch, J., Pitt, J.N., Reeves, N.L., Yao, J.C., Small, S., Desplan, C., et al. (2005). A major role for zygotic hunchback in patterning the *Nasonia* embryo. Development 132, 3705–3715.

58. Kumar, S., Stecher, G., and Tamura, K. (2016). MEGA7: Molecular Evolutionary Genetics Analysis Version 7.0 for Bigger Datasets. Mol Biol Evol 33, 1870–1874.

59. Guindon, S., Dufayard, J.F., Lefort, V., Anisimova, M., Hordijk, W., and Gascuel, O. (2010). New algorithms and methods to estimate maximum-likelihood phylogenies: assessing the performance of PhyML 3.0. Syst Biol 59, 307–321.

60. Huelsenbeck, J.P., and Ronquist, F. (2001). MRBAYES: Bayesian inference of phylogenetic trees. Bioinformatics 17, 754–755.

61. Letunic, I., and Bork, P. (2017). 20 years of the SMART protein domain annotation resource. Nucleic Acids Res.

62. Taylor, M.J., and Hoerauf, A. (1999). *Wolbachia* bacteria of filarial nematodes. Parasitol Today 15, 437–442.

63. McGraw, E.A., Merritt, D.J., Droller, J.N., and O’Neill, S.L. (2002). *Wolbachia* density and virulence attenuation after transfer into a novel host. Proc Natl Acad Sci U S A 99, 2918–2923.

64. Schindelin, J., Arganda-Carreras, I., Frise, E., Kaynig, V., Longair, M., Pietzsch, T., Preibisch, S., Rueden, C., Saalfeld, S., Schmid, B., et al. (2012). Fiji: an open-source platform for biological-image analysis. Nat Methods 9, 676–682.

