## Supplementary Materials for "The maternal effect gene *Wds* controls *Wolbachia* titer in *Nasonia*"

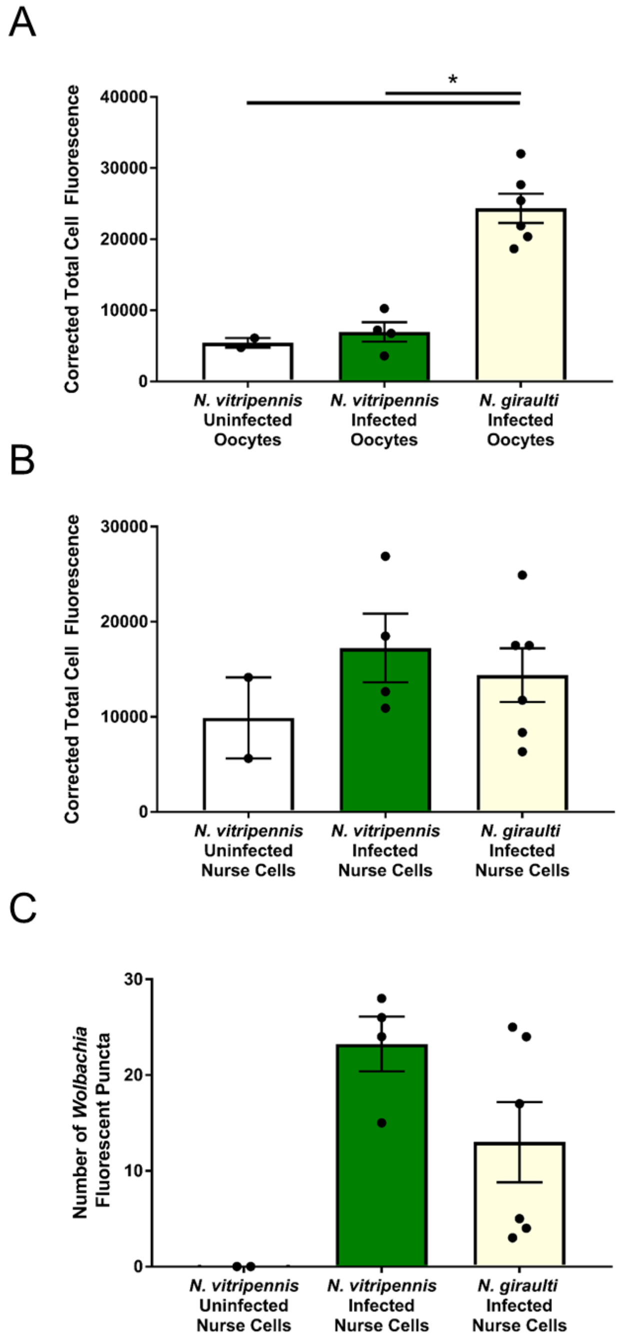


**Figure S1. Quantification of fluorescence in *Wolbachia*-infected *Nasonia* oocytes, Related to Figure 2.** (A) Bars represent the average corrected total cell fluorescence of *Wolbachia* in *N. vitripennis* 12.1 and *N. giraulti* IntG oocytes from stage 3 egg chambers. A *Wolbachia*-cured *N. vitripennis* line serves as a negative control. (B) Bars represent the average corrected total cell fluorescence of *Wolbachia* in *N. vitripennis* 12.1 and *N. giraulti* IntG nurse cell cytoplasms. A *Wolbachia*-cured *N. vitripennis* line serves as a negative control. (C) Bars represent counts of *Wolbachia* indicated fluorescent puncta in *N. vitripennis* 12.1 and *N. giraulti* IntG nurse cell cytoplasms. A *Wolbachia*-cured *N. vitripennis* line served as a negative control. * $p<0.05$, Kruskal Wallis test with Dunn's multiple correction. Error bars are mean ± S.E.M.

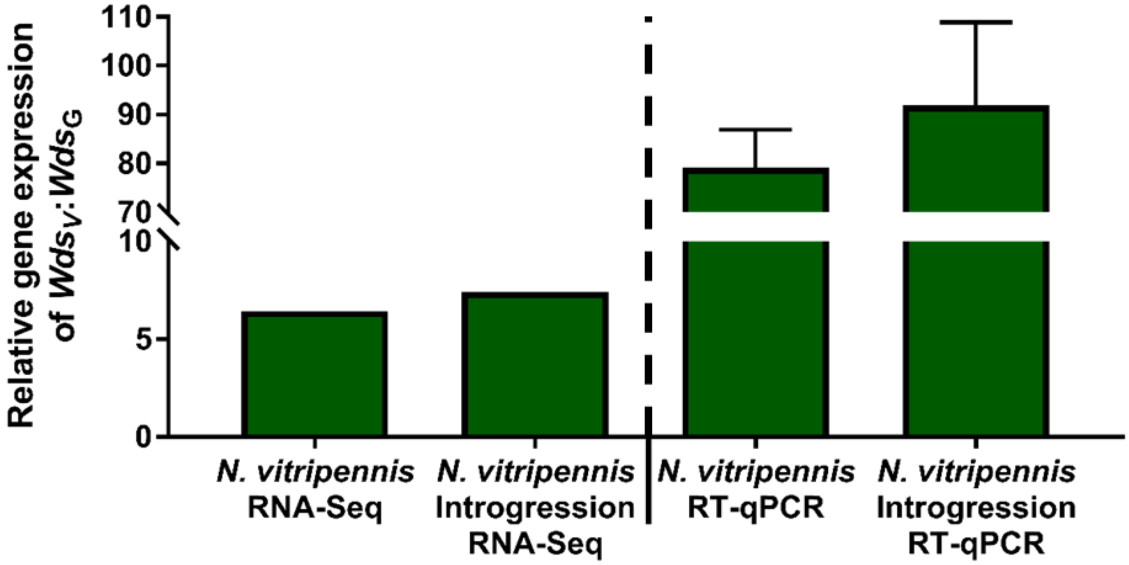


**Figure S2. RT-qPCR validation of *Wds$_v$* expression in ovaries of *N. vitripennis* and the *N. vitripennis* introgression IntC3 compared to *N. giraulti*, Related to Figure 5.** Bars represent the average fold change of *N. vitripennis* ovarian gene expression of *Wds$_v$* compared to *N. giraulti* IntG *Wds$_g$* expression. Error bars are mean ± S.E.M.

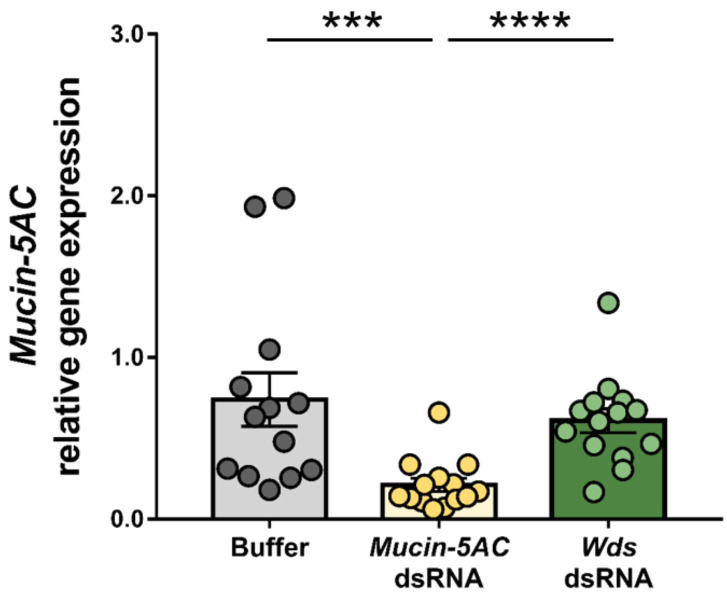


**Figure S3. Relative gene expression of *Mucin-5AC* in late pupae of dsRNA-injected females, Related to Figure 5.** Relative gene expression of *Mucin-5AC* in late pupae of *Mucin-5AC*-RNAi and *Wds*-RNAi females normalized to *Mucin-5AC* expression in buffer-injected females. ***p < 0.001, and ****p < 0.0001, Mann-Whitney U test. Error bars represent mean ± S.E.M.

**Table S1. Primers for *Nasonia* microsatellite markers, Related to Figure 3 and Figure 4.**

cM locations based on genetic linkage map from [S1]. N/A = sequence is absent in *N. giraulti* so no PCR product is generated. Primers were used for quantitative trait loci mapping (QTL), fine mapping in segmental introgression lines (FM), or both. All primers were designed as part of this study with the following exceptions: NvC1-20, Nv-20, and NvC3-18 are from [S2]; Nv184 is from [S3].

| Primer name | Chr | cM* | Primer Set (5' to 3') | Size *Nvit* (bp) | Size *Ngir* (bp) | Annealing Tm (ºC) | Used For |
|---|---|---|---|---|---|---|---|
| MM1.12 | 1 | 31.4 | F: GCGGTCCTGCTCCATTAACCGC<br>R: CCAGACTCGCGCGGGTGTATTT | 284 | 242 | 57 | QTL |
| MM1.13 | 1 | 32.9 | F: AGCTCCGAGAGCGCGAGTGA<br>R: TCCCGTGCCGACGCATACAC | 224 | 167 | 57 | QTL |
| MM1.14 | 1 | 35.8 | F: GCCGTCGAGAGACGAGCGAG<br>R: GCGCGGCTGGAGGATGCTTT | 219 | 266 | 57 | QTL |
| MM1.L521 | 1 | 38.7 | F: ACACGTCCCGATCCTTCTTTGAC<br>R: GCGCCTCACTTGTTGTGCAT | 118 | 160 | 54 | QTL |
| MM1.16 | 1 | 40.9 | F: ACGCGACTCCTTTCTCCGCA<br>R: GCGGAAATCGAATGCGCGGC | 233 | 199 | 56 | QTL |
| MM1.17 | 1 | 43.8 | F: TGCCTCGCGAGAGCGCAAAA<br>R: ACTGCTCTCGTCAAGGCCGC | 177 | 217 | 57 | QTL |
| NvC1-21 | 1 | 46.7 | F: GTAACAGTGAGATAAATGTG<br>R: TAGCAACGATAGTCCACG | 148 | N/A | 45 | QTL |
| MM1.057 | 1 | 49.6 | F: CTACCACATCTTTCGCCAGTTT<br>R: TCGAGTGATTAGAGATCGACGTT | 180 | 206 | 51 | QTL |
| MM1.L3567 | 1 | 53.3 | F: CGCTCTGTCTACCTGTCCCT<br>R: CGGCCACAAAGCAAATAGGC | 154 | 184 | 52 | QTL |
| MM1.31 | 1 | 56.2 | F: CGCATCATCAACCCCCGACCA<br>R: TCCGCGGCATAACCACTTGCT | 266 | 297 | 57 | QTL |
| MM1.32 | 1 | 57.7 | F: ACCGGGACGACTTGAGCGTA<br>R: ACAATGGGCGAATTTTTCTGCCG | 183 | 220 | 55 | QTL |
| MM2.13 | 2 | 19 | F: AAGACGAGAGCCGACGTTGC<br>R: GGCCTGCACGAGTGTGTATAGGG | 240 | 206 | 55 | QTL |
| MM2.15 | 2 | 21.9 | F:<br>TGGCAGATGACTCACGGAAATTAACAG<br>R:<br>CAGTTTTAGATGAGTTTATGAACTGTGT<br>C | 87 | 154 | 52 | QTL |
| MM2.17 | 2 | 24.8 | F: CGCCGACGTCGTTGCTGCTT<br>R: AGCTCCACAACGGCGGCATC | 143 | 99 | 58 | QTL |
| MM2.20 | 2 | 29.2 | F: TCTCCGTTAATTTCCAGCGCGT<br>R: TCTTCCAATCCACGGGAAAACTGGT | 207 | 168 | 55 | QTL |
| Nv-20 | 2 | 30.7 | F: TGACGAAGTATCCGAGAAG<br>R: TCGAAAAACGATATTGCTCG | 105 | 87 | 48 | QTL |
| MM2.26 | 2 | 32.9 | F: GCATCGCGTATGCTAATCTGCCG<br>R: GGCGGAGTGAGAGAGCGTTTCA | 220 | 172 | 56 | QTL |
| MM2.L5335 | 2 | 36.5 | F: CGCACGCGGTAATTGGCTTT<br>R: TGTCCACGGCTGCGATTTGT | 202 | 168 | 54 | QTL |
| MM2.28 | 2 | 38 | F: ACGCTTACACGCTGGTGAATGAA | 256 | 287 | 55 | QTL |

| | | | | | | | |
|---|---|---|---|---|---|---|---|
| | | | R: ACACCGTAATGCAATTTCCCGCT | | | | |
| MM2.30 | 2 | 40.9 | F: TGGATGCGAGCGCGGGTTAT<br>R: CCCATCGCTGATCCACGTTCTT | 135 | 172 | 55 | QTL |
| MM2.L6870 | 2 | 43.8 | F: GCTCTACACGGCGAAGGTCA<br>R: CGCGCTTCTCTTTATGCCCG | 140 | 191 | 54 | QTL |
| MM2.33 | 2 | 46 | F: ACGAAACTCTGTACTGTATACTCCGGT<br>R: CGGCGAGTCCTCGAGAGCAG | 204 | 250 | 55 | QTL |
| MM2.36 | 2 | 49.6 | F: GCCGTTGGAGAAATGTGCGGGA<br>R: TCGCGTATATTTTCCGTAGTCACGC | 178 | 139 | 55 | QTL |
| MM2.39 | 2 | 52.6 | F: ACCGTTACAAAGCGAGCGAGAAT<br>R: GCCGCCGCATAGCTCGATGA | 161 | 207 | 55 | QTL |
| MM2.40 | 2 | 54.8 | F: TCCGTTTATCGCGCTTCGGACG<br>R: CATCGGGCTGACCTTGGCCG | 179 | 211 | 57 | QTL |
| MM2.L7336 | 2 | 57.7 | F: CATTCATCGCTCGTGTGCGC<br>R: ACACATCTCTCCGAACGGCG | 118 | 85 | 54 | QTL |
| MM2.44 | 2 | 60.6 | F: TCGACGGAAGCGAGGACGAG<br>R: CTGGGCCGCAACGGTAAGCA | 203 | 172 | 56 | QTL |
| MM2.49 | 2 | 68.6 | F: ACTGTTGCAGATGATGATGGTAATTT<br>R: TCTGAAACATGCAACAATCAGGT | 146 | 92 | 51 | QTL |
| MM3.14 | 3 | 17.5 | F: CTCTCGAAGCCGCGCGTGAA<br>R: AGCCAGCTTTGCTTTCGACCG | 231 | 206 | 56 | QTL |
| MM3.15 | 3 | 20.4 | F: ACACACGTTGTGCGGGGGTG<br>R: GGTCGAAAATTTCTGCGCAGCCT | 106 | 152 | 56 | QTL |
| MM3.17 | 3 | 23.4 | F: TGCGCGATGGCTGCTGTGAT<br>R: TCGAGCGCAATAAACGCCGC | 126 | 170 | 57 | QTL |
| MM3.19 | 3 | 26.3 | F: GCGGAAATTCTCGCCCCTGC<br>R: TCCCATCATCAAAACGAAAAAGTCGC | 177 | 220 | 55 | QTL, FM |
| MM3.22 | 3 | 29.2 | F: TCTCCTCCTGCTTCGGCCCC<br>R: TCGTTCATCGTTCGTCATCGCA | 116 | 146 | 55 | QTL, FM |
| MM3.L8514 | 3 | 32.1 | F3: GCGGCGAGAAGAACAGAAGCGA<br>R3: ACGTTGTCGCTGTTCTTCTTGCAGT | 488 | 279 | 57 | FM |
| MM3.23 | 3 | 32.9 | F: TTGAAGGGCTCATGGTCGCA<br>R: CGCGAAACAGCGCACACG | 183 | 219 | 55 | QTL, FM |
| MM3.L8610 | 3 | 34.3 | F1: CGCGTATGCTTGATGTCGGC<br>R1: AACACAGAGGAATATGCGGGA | 172 | 129 | 51 | FM |
| MM3.L8623 | 3 | 34.3 | F: TTGGAGTTTCGCACAAGAGC<br>R: CTACCGCCGAGAAAAGTGC | 185 | 136 | 54 | FM |
| MM3.L8651 | 3 | 35.0 | F: AGATGAGAAGAAAGAGGAAGCCC<br>R: ATGGCGATTTCTTCATGTCGC | 164 | 117 | 54 | FM |
| MM3.L8678 | 3 | 35.0 | F: GCAGCCAGGGAGTGATATGCT<br>R: AAAGGCCGACGACGAGAGAC | 186 | 138 | 54 | QTL, FM |
| MM3.L8716 | 3 | 36.5 | F: TGCCCATCAAAGGTGAGAGG<br>R: CGGACTCACTGTTTGCCCAG | 237 | 208 | 54 | FM |
| MM3.L8724 | 3 | 36.5 | F: CCTCTGTCTGTGCTTTTACGG<br>R: TTCCCGTAAACACGATTGCC | 105 | 82 | 54 | FM |
| MM3.L8756 | 3 | 36.5 | F: CGCGTGTCGTGTGGACGTAA<br>R: TCAAACATCCGCGAGAGTCGA | 115 | 157 | 54 | FM |

| | | | | | | | |
|---|---|---|---|---|---|---|---|
| MM3.L8780 | 3 | 37.2 | F1: TTAACCGAATAGCACCGCCG<br>R1: AAATCCCAGATCCCGCACTAG | 185 | 140 | 51 | FM |
| MM3.L8790 | 2 | 37.2 | F1: ATTTACCGACGCGCAACAGC<br>R1: AGGGCGGAGAGATTAGATTTCCC | 159 | 195 | 53 | FM |
| MM3.L8813 | 3 | 37.2 | F: CCGAGTGTGGGAGGTTTGACA<br>R: TGTCAGCCGAGAATAGGCCG | 177 | 148 | 54 | FM |
| MM3.L8850 | 3 | 37.2 | F: TGGTTGAGAGATCCACGCGA<br>R: TCCGCGTTTACAACCAACATGG | 159 | 206 | 53 | FM |
| NvC3-18 | 3 | 38 | F: GCCCAAATCATGCTTTCG<br>R: GTTGTTCTTAAATGTGTATTCC | 104 | N/A | 48 | QTL |
| MM3.29 | 3 | 39.4 | F: GGCCGATTTTCTCGACAGACC<br>R: GCGAGGGAGAGCGAACGTC | 241 | 285 | 53 | QTL, FM |
| MM3.L10131 | 3 | 40.2 | F: TGATGCGTTCTCGCCTTTCC<br>R: CGACCGCAGAGCAACGATCA | 155 | 204 | 53 | FM |
| MM3.L10212 | 3 | 41.6 | F: CCTCCCAAATCACTTCCGCGT<br>R: TCAGCGCAATCGTTACCCTT | 108 | 135 | 52 | QTL, FM |
| Nv184 | 3 | 44.5 | F: GCGTCATCGATGCATTTCTT<br>R: TCTCGGGAGAGATTCAGTACG | 209 | 141 | 49 | QTL |
| MM3.L10340 | 3 | 45.3 | F: CGAAACACCATTCGCAACGAGT<br>R: TGTCGCATCGAGAACTGCA | 194 | 167 | 52 | FM |
| MM3.29.7M | 3 | 45.3 | F:<br>CCAGTTGGATAATTCTTGAGGTCTTTC<br>R: ACTTTGCTTGGCCCGACGAT | 148 | 118 | 52 | FM |
| MM3.35 | 3 | 46.7 | F: GTACGTGAACCGGAAGTGTTT<br>R: GACGGCTGCTACCGGCTATA | 111 | 161 | 51 | QTL, FM |
| MM3.36 | 3 | 48.2 | F: ATTCGCGCCGCGGCTAATGG<br>R: TTCCATACGTGTGGCAGGCG | 150 | 197 | 55 | FM |
| MM3.37 | 3 | 50.4 | F: ACAAGCTTCGCACACACCGCA<br>R: CGGTCGAAGAAGCGTCGCACA | 185 | 157 | 58 | QTL, FM |
| MM3.L10502 | 3 | 54.8 | F: GCGCGAAACGACGAGGAATT<br>R: CGAGCGTCGTGTGCTCTTCT | 63 | 94 | 54 | QTL |
| MM3.41 | 3 | 58.4 | F: ACCGTGGGTCCGTGCAAC<br>R:<br>GGTTTGTACTTCATCGTGAGGCAATCG | 186 | 142 | 55 | QTL, FM |
| MM3.L10553 | 3 | 63.5 | F: GCGCTTAATTGCGTCGTGTT<br>R: CCGGTGCGGTTTCTTCTCCT | 196 | 234 | 52 | QTL |
| MM3.43 | 3 | 65.7 | F:<br>CGGCTGTTTATATTCCTCACCTGACGC<br>R: GCAGCGACGAATCAGGAAATGCG | 138 | 158 | 57 | QTL |
| MM3.45 | 3 | 69.4 | F: CGATTATGCAAACGACGCGA<br>R: TTCCGATCACGATTCTCTCCTT | 222 | 168 | 51 | QTL |
| MM3.L10661 | 3 | 73 | F: CCCTCCGATTATAGATGCAAGTGTCA<br>R: GGCAGTAGTGGCTCTCTTTGCT | 159 | 181 | 54 | QTL |

**Table S2. Mapping statistics for RNA-seq of *Nasonia* ovaries, Related to Figure 5.**

Nvit: *N. vitripennis* strain 12.1; IntG: *N. giraulti* strain IntG; IntC3: *N. giraulti* strain IntG introgressed with a chromosome 3 region from *N. vitripennis* strain 12.1. Analysis of the RNA-seq paired-end reads was performed in CLC Genomics Workbench 8.

| Sample | # of Reads after QC | # of Mapped Paired Reads | % of Total Paired Reads Mapped | # of Mapped Intergenic Gene Reads | # of Mapped Gene Reads | # of Uniquely Mapped Gene Reads |
|---|---|---|---|---|---|---|
| Nvit-1 | 30,390,144 | 22,202,334 | 73.06 | 1,030,126 | 10,071,041 | 9,635,225 |
| Nvit-2 | 34,665,838 | 25,567,636 | 73.75 | 1,253,382 | 11,530,436 | 11,022,442 |
| Nvit-3 | 30,525,996 | 23,230,600 | 76.10 | 1,130,430 | 10,484,870 | 10,028,047 |
| Nvit-4 | 26,369,828 | 19,186,084 | 72.76 | 1,046,799 | 8,546,243 | 8,152,609 |
| IntC3-1 | 44,261,706 | 33,917,290 | 76.63 | 1,293,427 | 15,665,218 | 15,064,507 |
| IntC3-2 | 49,692,100 | 35,712,884 | 71.84 | 1,658,981 | 16,197,461 | 15,517,397 |
| IntC3-3 | 34,959,996 | 25,770,826 | 73.72 | 1,180,170 | 11,705,243 | 11,202,164 |
| IntC3-4 | 42,765,380 | 30,299,978 | 70.85 | 1,327,688 | 13,822,301 | 13,261,676 |
| IntG-1 | 36,821,016 | 27,095,510 | 73.59 | 1,051,834 | 12,495,921 | 12,020,977 |
| IntG-2 | 35,451,744 | 26,505,936 | 74.77 | 1,114,883 | 12,138,085 | 11,647,781 |
| IntG-3 | 40,863,430 | 31,432,456 | 76.92 | 1,234,399 | 14,481,829 | 13,913,515 |
| IntG-4 | 38,523,210 | 25,325,636 | 65.74 | 1,744,894 | 10,917,924 | 10,470,342 |
| IntG-5 | 38,697,378 | 27,021,558 | 69.83 | 1,201,004 | 12,309,775 | 11,765,553 |

**Table S3. Significant differentially expressed genes in R6-3 candidate region, Related to Figure 5.**

Mean number of reads for each gene was calculated by dividing the total number of reads from all replicates that mapped to the gene by the number of replicates (N = 4-5 for each *Nasonia* strain). Positive fold change = upregulation in *N. vitripennis* 12.1 or introgression line IntC3, while negative fold change = upregulation in *N. giraulti* IntG. Fold change and p-values were calculated using EdgeR in CLC Genomics.

| NCBI Gene ID | NCBI Annotated Gene Name | Mean Reads for Nvit | Mean Reads for IntC3 | Mean Reads for IntG | Fold Change (Nvit/IntG) | p-value (FDR-corrected) | Fold Change (IntC3/ IntG) | p-value (FDR-corrected) |
|---|---|---|---|---|---|---|---|---|
| LOC100679092 | Uncharacterized (*Wds*) | 38.8 | 66.3 | 7.60 | 6.44 | 3.09E-10 | 7.42 | 2.07E-14 |
| LOC100118928 | Dentin Sialophosphoprot ein-like | 23.5 | 18 | 11.2 | 2.67 | 2.98E-3 | 1.40 | 1 |
| LOC100118450 | Abnormal long morphology protein 1-like | 81.3 | 117 | 234 | -2.26 | 2.29E-8 | -2.30 | 5.32E-7 |
| LOC100679834 | Myb-like protein | 147.5 | 380 | 546 | -2.89 | 2.76E-5 | -1.63 | 3.35E-3 |
| LOC100114497 | Girdin | 640.5 | 1281 | 2508 | -3.05 | 1.46E-40 | -2.24 | 7.42E-19 |
| LOC103317608 | Interaptin-like | 144 | 140 | 517 | -2.82 | 3.48E-16 | -4.25 | 1.28E-27 |
| LOC100678491 | Uncharacterized | 26 | 46.3 | 152 | -4.49 | 5.25E-15 | -3.76 | 2.43E-13 |

**Table S4. Blastp homology to Wds$_V$ protein sequence, Related to Figure 6.**

BLASTp analysis performed on uncharacterized protein LOC100679092 (NCBI accession number XP_008213336.1) using the non-redundant protein sequences (nr) database.

| BLASTp Result Organisms | Accession | BLASTp E-value | Query coverage | % identity | Reciprocal BLASTp E-value to LOC100679092 |
|---|---|---|---|---|---|
| **Hymenopterans** | | | | | |
| Uncharacterized protein LOC100679092 [*Nasonia vitripennis*] | XP_008213336.1 | 1.00E-143 | 100% | 100% | 1.00E-143 |
| hypothetical protein TSAR_005991 [Trichomalopsis sarcophagae] | OXU27029.1 | 3.00E-99 | 100% | 95% | 1.00E-101 |
| Uncharacterized protein LOC106659966 [Trichogramma pretiosum | XP_014238261.1 | 4.00E-24 | 74% | 42% | 1.00E-21 |
| Uncharacterized protein LOC106636480 [Copidosoma floridanum] | XP_014204363.1 | 5.00E-18 | 98% | 35% | 2.00E-17 |
| Uncharacterized protein LOC106793189 [Polistes canadensis] | XP_014615392.1 | 2.00E-09 | 78% | 29% | 5.00E-07 |
| Uncharacterized protein LOC100878703 [Megachile rotundata] | XP_003702181.1 | 6.00E-08 | 56% | 32% | 4.00E-05 |
| Uncharacterized protein LOC107264954 isoform X2 [Cephus cinctus] | XP_015589287.1 | 5.00E-08 | 30% | 55% | 3.00E-08 |
| Uncharacterized protein LOC107186552 [Dufourea novaeangliae] | XP_015429933.1 | 3.00E-07 | 56% | 31% | 2.00E-03 |
| Uncharacterized protein LOC105276036 [Ooceraea biroi] | XP_011331664.1 | 4.00E-07 | 91% | 31% | 5.10E-02 |
| Uncharacterized protein LOC108574276 [Habropoda laboriosa] | XP_017792325.1 | 2.00E-06 | 75% | 26% | 1.00E-03 |
| Uncharacterized protein LOC100650078 [Bombus terrestris] | XP_003396525.1 | 2.00E-06 | 56% | 31% | 8.00E-05 |
| Uncharacterized protein LOC107963970 [Apis mellifera] | XP_001121053.1 | 2.00E-05 | 56% | 31% | 1.00E-02 |
| Uncharacterized protein LOC109860277 [Pseudomyrmex gracilis] | XP_020294850.1 | 3.00E-04 | 68% | 28% | 4.00E-03 |
| **Dipterans** | | | | | |
| Uncharacterized protein LOC109418023 [Aedes albopictus] | XP_019547726.1 | 1.50E-02 | 32% | 38% | 1.60E+00 |
| GL10758 [Drosophila persimilis] | XP_002015939.1 | 1.90E-02 | 32% | 36% | 8.90E-02 |
| Uncharacterized protein Dpse_GA24472 [Drosophila pseudoobscura] | XP_002138301.1 | 7.60E-02 | 32% | 34% | 8.60E-02 |

**Table S5. Primers for *Nasonia* gene sequencing and allelic genotyping, Related to Figure 4.**
cM locations based on genetic linkage map from [S1].

| Primer Name | Chr | cM* | Primer Set (5' to 3') | Annealing Temp (ºC) | Product Size (bp) |
|---|---|---|---|---|---|
| LOC100119494 | 3 | 35.0 | F: CGCACTCGCACAACATTCCC<br>R: GCCTTGCTCTTCTCCTTCTCCG | 57 | 617 |
| Mucin-5AC seq | 3 | 35.0 | F: TCGGCAAGAAGATGGGCGTC<br>R: GCGTCGTTGTGGTGGTTGTG | 57 | 747 |
| LOC100679324 | 3 | 36.5 | F: TATTGCCCCTCGCCCCATTG<br>R: CTGGTTCATAGCGTTGTTTGACATC | 56 | 715 |
| LOC100119358 | 3 | 36.5 | F: CATCATCGCCCTCGTCCTCTTC<br>R: ACCGTCTGCGTCACTTCCTG | 56 | 682 |
| LOC100119295 | 3 | 36.5 | F: CCAGGACCCAGACCAGGATTAG<br>R: AACCCACTTCTACCAGCCCC | 55 | 670 |
| LOC100119259 | 3 | 36.5 | F: TTGACCACACCGACAACAAC<br>R: GCATTTCATAAGTTCCGCCAGAG | 53 | 670 |
| LOC100679834 | 3 | 36.5 | F: GCCGATTACTGGACCGACAG<br>R: GTTGGGGTTGCGGATAGTTCG | 54 | 607 |
| LOC103317434 | 3 | 36.5 | F: ACGGGTATTTCAGCCTTCGCC<br>R: ACAACTCACACCTTCCCACCG | 57 | 723 |
| *Wds* seq | 3 | 36.5 | F: GTTCCTGATACTGCTCGCCG<br>R: ACTTTGCTTGGCCCGACGAT | 55 | 250 |
| LOC100118928 | 3 | 36.5 | F: CGAGCGAAGCACCGAGTTAC<br>R: GCAGGCGACAGTTCTCAACG | 55 | 677 |
| LOC100679277 | 3 | 36.5 | F: TTCGGGTCTTTTGTTATTGCGAG<br>R: TATCCTCCGTGTCCTCCGTG | 53 | 616 |
| LOC100118712 | 3 | 36.5 | F: TAGACCACGAACGCAACCTCG<br>R: GCTTCCCAAGAACCCATCCC | 57 | 632 |
| LOC100118529 | 3 | 36.5 | F: GTCTCGGCGGGTTTTGATGG<br>R: GCGTCCTTTGGTGGCTGTTG | 55 | 685 |
| LOC100118450 | 3 | 36.5 | F: ATGGAAAAGGCATCGGTAAGCG<br>R: GCAACTCAGAAATCGTCCTGCG | 55 | 630 |
| LOC100114497 | 3 | 36.5 | F: ACGAGTCATCTTCTATGGTTTTGGC<br>R: TGTGGCAGGCGTTTGAGTATC | 54 | 375 |
| LOC100678491 | 3 | 36.5 | F: GGATCACAACCAAAAGTTCCTG<br>R: GGTACGGCCTAAACACGG | 54 | 540 |

**Table S6: Primers for dsRNA constructs and RT-qPCR of *Nasonia* genes. Related to Figure 5.**

| NCBI Gene ID | Gene Name | Name of Primer Set | Primer sequences (5' to 3') | Product Size (bp) |
|---|---|---|---|---|
| LOC100679092 (*Nasonia*) | Uncharacterized | Unchar RNAi | F: CGTTCCTGATACTGCTCGCC<br>R: CCACCTGTTGCCTGTAGACG | 438 |
| | | Unchar1 qPCR | F: ACCTACTGCTGACATCGTTCC<br>R: AGCCCGTCTCTTGTTTCACG | 165 |
| LOC100679394 (*Nasonia*) | Mucin-5AC | Mucin5ac RNAi | F: TCGGCAAGAAGATGGGCGTC<br>R: GCGTCGTTGTGGTGGTTGTG | 627 |
| | | Mucin5ac qPCR | F: AAGGCTCGTGGAAGACTGCG<br>R: TGGCGGCGTCCTGTTGTATC | 145 |
| malE (*E. coli*) | Maltose transporter subunit | MalE RNAi | F: ATTGCTGCTGACGGGGGTTAT<br>R: ATGTTCGGCATGATTTCACCTTT | 495 |
| LOC100115795 (*Nasonia*) | 60S Ribosomal protein L32 | RP49 qPCR | F: CAAGCGTAACTGGAGGAAGC<br>R: CTGCTAACTCCATGGGCAAT | 221 |

**Supplemental References**

S1. Desjardins, C.A., Gadau, J., Lopez, J.A., Niehuis, O., Avery, A.R., Loehlin, D.W., Richards, S., Colbourne, J.K., and Werren, J.H. (2013). Fine-scale mapping of the *Nasonia* genome to chromosomes using a high-density genotyping microarray. G3 *3*, 205-215.

S2. Rutten, K.B., Pietsch, C., Olek, K., Neusser, M., Beukeboom, L.W., and Gadau, J. (2004). Chromosomal anchoring of linkage groups and identification of wing size QTL using markers and FISH probes derived from microdissected chromosomes in *Nasonia* (Pteromalidae: Hymenoptera). Cytogenet Genome Res *105*, 126-133.

S3. Beukeboom, L.W., Niehuis, O., Pannebakker, B.A., Koevoets, T., Gibson, J.D., Shuker, D.M., van de Zande, L., and Gadau, J. (2010). A comparison of recombination frequencies in intraspecific versus interspecific mapping populations of *Nasonia*. Heredity (Edinb) *104*, 302-309.
