## Supplementary Materials for "The maternal effect gene *Wds* controls *Wolbachia* titer in *Nasonia*"

**KEY RESOURCES TABLE**

| REAGENT or RESOURCE | SOURCE | | IDENTIFIER |
| --- | --- | --- | --- |
| Antibodies | | | |
| Mouse monoclonal Anti-human Hsp60 | Sigma | Cat#H3524 | |
| Goat anti-mouse IgG Alexa Fluor 594 | Thermo Fisher Scientific | Cat#A-11005 | |
| Chemicals, Peptides, and Recombinant Proteins | | | |
| GelRed | Biotium | | Cat#41003-1 |
| SYTOX Green Nucleic Acid Stain | Thermo Fisher Scientific | | Cat#S7020 |
| ProLong Gold Antifade Mountant | Thermo Fisher Scientific | | Cat#P36930 |
| ProLong Diamond Antifade Mountant | Thermo Fisher Scientific | | Cat#P36970 |
| Critical Commercial Assays | | | |
| Gentra Puregene Tissue Kit | Qiagen | | Cat#158667 |
| DNeasy Blood and Tissue Kit | Qiagen | | Cat#69504 |
| Direct-zol RNA Miniprep kit | Zymo Research | | Cat#R2050 |
| Nucleospin RNA/Protein Kit | Macherey-Nagel | | Cat#740933.50 |
| iQ SYBR Green Supermix | Bio-Rad | | Cat#1708882 |
| iTaq Universal SYBR Green Supermix | Bio-Rad | | Cat#1725122 |
| GoTaq Green Master Mix | Promega | | Cat#M7123 |
| REPLI-g Mini Kit | Qiagen | | Cat#150023 |
| QIAquick PCR Purification Kit | Qiagen | | Cat#28104 |
| QIAquick Gel Extraction Kit | Qiagen | | Cat#28704 |
| RQ1 RNase-free DNase | Promega | | Cat#M6101 |
| DNA-free DNA Removal Kit | Thermo Fisher Scientific | | Cat#AM1906 |
| Qubit RNA HS Assay Kit | Thermo Fisher Scientific | | Cat#Q32852 |
| Qubit dsDNA Broad Range Assay Kit | Thermo Fisher Scientific | | Cat#Q32850 |
| SuperScript VILO cDNA Synthesis Kit | Thermo Fisher Scientific | | Cat#11754050 |
| MEGAScript RNAi kit | Thermo Fisher Scientific | | Cat#AM1626 |
| Deposited Data | | | |
| RNA sequencing reads | This paper | | SRA: PRJNA430433 |
| Experimental Models: Organisms/Strains | | | |
| *Nasonia vitripennis* 12.1 | Perrot-Minnot and Werren, 1999 | | N/A |
| *Nasonia giraulti* IntG12.1 | Chafee et al., 2011 | | N/A |
| *Wolbachia* groEL (qPCR forward primer): CAACCTTTACTTCCTATTCTTG | Bordenstein et al., 2006 | | N/A |
| *Wolbachia* groEL (qPCR reverse primer): CTAAAGTGCTTAATGCTTCACCTTC | Bordenstein et al., 2006 | | N/A |
| *Nasonia* S6K (qPCR forward primer): GGCATTATCTACAGAGATTTGAAACCAG | Bordenstein and Bordenstein, 2011 | | N/A |
| *Nasonia* S6K (qPCR reverse primer): CAAAGCTATATGACCTTCTGTATCAAG | Bordenstein and Bordenstein, 2011 | | N/A |
| Primers for *Nasonia* microsatellite markers, see Table S1 | This paper | | N/A |
| Primers for *Nasonia* gene sequencing, see Table S5 | This paper | | N/A |
| Primers for dsRNA construct and RT-qPCR, see Table S6 | This paper | | N/A |
| Software and Algorithms | | | |
| Geneious Pro | Biomatters | | <http://www.geneious.com> |
| MATLAB | MathWorks | | <https://www.mathworks.com/products/matlab.html> |
| R Software | R Project | | https://www.r-project.org/ |
| R package: *R/qtl* | Broman et al., 2003 | | http://www.rqtl.org/ |
| CLC Genomics Workbench | Qiagen | | https://www.qiagenbioinformatics.com/products/clc-genomics-workbench/ |
| MEGA7 | Kumar et al., 2016 | | http://www.megasoftware.net/ |
| SWAKK Bioinformatics Server | Liang et al., 2006 | | http://ibl.mdanderson.org/swakk/ |
| SMART Online Software | Letunic and Bork, 2017 | | <http://smart.embl-heidelberg.de> |
| FIJI (ImageJ) | Schindelin et al., 2012 | | https://fiji.sc/ |
| GraphPad Prism | GraphPad Software | | https://www.graphpad.com/scientific-software/prism/ |
